# α-Synuclein pathology and reduced neurogenesis in the olfactory system affect olfaction in a mouse model of Parkinson’s disease

**DOI:** 10.1101/2022.08.19.504562

**Authors:** Eduardo Martin-Lopez, D.J. Vidyadhara, Teresa Liberia, Sarah J. Meller, Leah E. Harmon, Ryan M. Hsu, Kimberly Han, Betül Yücel, Sreeganga S. Chandra, Charles A. Greer

**Affiliations:** Department of Neurosurgery, Yale University School of Medicine, 310 Cedar Street, New Haven, CT 06510, USA; Department of Neurology, Yale University School of Medicine, 310 Cedar Street, New Haven, CT 06510, USA; Department of Neuroscience, Yale University School of Medicine, 310 Cedar Street, New Haven, CT 06510, USA

**Keywords:** α-Synuclein, A30P, Parkinson’s disease, olfaction, neurogenesis, endocytosis, presynaptic proteins

## Abstract

Parkinson’s Disease (PD) is characterized by multiple symptoms including olfactory dysfunction, whose underlying mechanisms remain unclear. Here, we explored pathological changes in the olfactory pathway of transgenic (Tg) mice expressing the human A30P mutant α-synuclein (α-syn) (α-syn-Tg mice) at 6-7 and 12-14 months of age, representing early and late-stages of motor progression, respectively. α-Syn-Tg mice at late stages exhibited olfactory behavioral deficits, which correlated with severe α-syn pathology in projection neurons of the olfactory pathway. In parallel, olfactory bulb (OB) neurogenesis in α-syn-Tg mice was reduced in the OB granule cells at 6-7 months, and OB periglomerular cells at 12-14 months, respectively, both of which could contribute to olfactory dysfunction. Proteomic analyses showed a disruption in endo- and exocytic pathways in the OB during the early stages which appeared exacerbated at the synaptic terminals when the mice developed olfactory deficits at 12-14 months. Our data suggest that, 1) the α-syn-Tg mice recapitulate the olfactory functional deficits seen in PD; 2) olfactory structures exhibit spatiotemporal disparities for vulnerability to α-syn pathology; 3) α-syn pathology is restricted to projection neurons in the olfactory pathway; 4) neurogenesis in adult α-syn-Tg mice is reduced in the OB; and 5) synaptic endo- and exocytosis defects in the OB may further explain olfactory deficits.

## Introduction

Parkinson’s Disease (PD) affects quality of life due to disparate motor and non-motor clinical symptoms. A common non-motor symptom in PD is hyposmia or anosmia which is seen in 75-90% of affected individuals (1–5). Due to its high prevalence, anosmia is now included as a diagnostic criterion for PD by the Movement Disorder Society (6). Yet, the mechanisms underlying olfactory dysfunction in PD remain unclear and not fully investigated.

PD is pathologically characterized by the neuronal accumulation of insoluble, cytoplasmic protein inclusions known as Lewy bodies (LBs) and Lewy neurites (7–9). LBs consist mainly of α-synuclein (α-syn), a synaptic protein that regulates synaptic vesicle exo- and endocytosis (10, 11). Thus, the formation of LBs is associated with synaptic dysfunction and neurodegeneration (12, 13). In PD, pathological α-syn can assemble into multimers that form neurotoxic oligomers and fibrils (14, 15), whose accumulation is accompanied by phosphorylation of the protein at serine 129 (pSer129-α-syn) (16, 17) that increases with disease progression (18, 19). Indeed, ~90% of α-syn in LBs is phosphorylated at this position (20, 21) and pSer129-α-syn antibodies are used to identify α-syn pathology in animal models of PD (22). Furthermore, increased levels of pSer129-α-syn are found in postmortem human olfactory bulbs (23).

The events triggering LB pathology in the brain remain speculative, but there is evidence suggesting that α-syn misfolding and aggregation may originate in the olfactory epithelium and gastric system, after which the disease propagates to the brain via olfactory and enteric pathways (24–29). Although environmental factors are reported to be involved in α-syn misfolding and PD pathogenesis (30, 31), other evidence points to genetic factors including mutations in *GBA, LRRK2* and *SNCA* genes as a cause of PD and associated LB pathology (30, 32, 33). Among these, point mutations in the gene *SNCA* encoding for α-syn (e.g., A30P, A53T, E46K, G51D and H50Q), generate aggressive, fully penetrant, early-onset forms of PD (34–38). These and other mutations have been used to generate animal models to study familial forms of PD (39, 40).

As noted above, anosmia is among the criteria used to diagnose PD. In a comparable vein, analyses of mouse models with *SNCA* mutations suggested the development of olfactory deficits (41–44). To understand more fully the mechanisms that may account for any olfactory deficits associated with *SNCA* mutations, in this study we used a mouse model that overexpressed human A30P mutant α-syn (45) (α-syn-Tg mice) to investigate the olfactory system. Our findings indicated a preferential accumulation of α-syn pathology in projection neurons throughout the olfactory pathway and a decreased OB neurogenesis. In addition, we detected a decrease in synaptic exo- and endocytic protein expression, events that were likely to account for the olfactory dysfunction in the α-syn-Tg mice.

## Materials and Methods

### Animals

All experiments were performed in C57BL6/J congenic Thy-1 human A30P mutant α-syn mice (α-syn-Tg) and their wild-type (WT) control littermates (45). Analyses were carried out on either 6–7 months (early stage) or 12–14 months (late stage) cohorts of mice. Mice were raised and housed at Yale University vivarium on a 12 h light-dark cycle with access to standard chow *ad libitum*. All protocols were approved by Yale University Institutional Animal Care and Use Committee.

### Buried food tests

To detect behavioral olfactory deficits, we conducted two standardized buried food tests to optimize the detection of any anomalies in olfactory function in the early vs. late stages (46–48). Both tests determine the ability of mice to detect a food reward hidden in the bedding of clean housing cages.

On 6-7 months mice (WT: n=7, 3 males and 4 females; α-syn-Tg: n=7, 5 males and 2 females) testing took place over 3 consecutive days(49). On day 1 (*habituation day*) mice were placed individually in clean polycarbonate cages (Acecaging, USA) filled with 3 cm of 1/8” corncob bedding (Teklad, 7092). A piece of cheese flavored cookie (Better Cheddars, Nabisco) was left on the surface of the bedding and mice were allowed to habituate to the cookie smell for 5 min (*odor familiarization*). Before day 2, food consumption was restricted by providing them with only 1.82 g of standard lab pellets (7 Kcal/mouse). On day 2 (*pre-test partial food deprivation test*), a piece of cheese cookie was hidden inside the bedding along the perimeter of the testing cage and the latency (in seconds) to find the cookie was recorded (Trial 1). A latency longer than 300 sec. was considered “failed” and mice were scored with 300 sec. Four identical trials were conducted with an interval of 1 h between trials. The bedding of each cage was changed between trials and the cookie was buried at different locations between days and trials. At the end of day 2 mice were fully deprived of food for 24 h, and on day 3, *score latency olfactory tests* were conducted as described for day 2. Weights were monitored daily due to food restrictions and performance between WT and α-syn-Tg males and females was compared to evaluate the influence of sex in the olfactory performance. The latency scores for each animal were calculated as the average between the trials and testing days. To exclude a bias produced by lack of motivation for food or because of motor deficits, a surface food test was also performed as previously described (50). Briefly, a piece of the same cheese cookie was placed in a specific location of the surface of each cage with clean bedding, and mice were dropped in the opposite side of the cage. The latency to find and showing interest for the food was recorded for each individual animal.

Mice at 12-14 months (WT: n=6, 4 males and 2 females; α-syn-Tg: n=8 males) were tested using an equivalent protocol (51). Tests were carried out over 3 consecutive days in clean polysulfone cages (Techniplast) using a piece of sweetened cereal. As described for the 6-7 months mice, on day 1 (*odor familiarization*), the 12-14 months mice were exposed to a piece of cereal (Chocolate Cheerios, General Mills) placed in each cage. The cereal was left for consumption overnight and only those mice that consumed the cereal were selected for the further analysis. On day 2 mice were deprived of food for 24 h. On day 3 mice were habituated to the testing environment by placing them in a clean cage with 3 cm of clean 1/8” corncob bedding (Teklad, 7092) for 60 min., after which mice were returned to home cage. Test trials (*score latency olfactory test*) were as follows: during <u>Trial 1</u> a piece of cereal was buried in a selected location in the test cage and the mouse then returned to the test cage at a location opposite the site of the buried cereal. The latency to find the cereal was recorded; mice that took longer than 300 sec. to find the cereal were scored as 300 sec. <u>Trials 2</u> and <u>3</u> were performed identically but the pieces of cereal were placed in a different corner of the cage. The latency scores for each animal were the average for the 3 trials. For each animal, this test was repeated for 3 weeks, and the scores were averaged. As for the 6-7 months mice, body weights were monitored to ensure the mice were in good health and a surface food test as described above was also carried out to exclude bias due to lack of motivation for the food or motor deficits.

### Tissue processing and immunohistochemistry

To obtain tissues for immunohistochemistry (IHC), mice were deeply anesthetized with an overdose of Euthasol (Virbac) and transcardially perfused first with 5 mL saline solution, followed by 1 mL/g of body weight of ice-cooled 4% paraformaldehyde (PFA) in PBS. Fixed brains were removed from skulls and postfixed for 24 h at 4°C. The brains were then transferred to 30% sucrose-PBS for cryoprotection and embedded in Tissue-Tek OCT compound (Fisher Scientific, Cat# 4585) for sectioning. Sections 25 μm thick were obtained using a Reichert Frigocut cryostat (E-2800) in the coronal plane, then air dried and stored at −80°C until use.

Before staining, sections were thawed in a slide warmer at 60°C for 10 min. and rehydrated in PBS for 10 min. An antigen unmasking step was conducted by incubating sections in 0.01M citrate buffer (pH 6.0) at 65°C for 35 min., followed by 5 min. incubation in ice-cooled citrate buffer to perform a heat-shock. Then, slides were washed three times, 10 min. each with PBS + 0.1% Triton X-100 (Millipore-Sigma, Cat# x100) (PBST), and sections were incubated with blocking solution made of PBST supplemented with 5% normal goat serum (NGS) (Accurate Chemicals, Cat# CL1200) and 0.01% of protease-free bovine serum albumin (BSA) (Sigma-Aldrich, Cat# A3059) for 1 h at room temperature (RT). The blocking solution was removed, and sections were incubated overnight with primary antibodies (Table 1) diluted in 10% diluted blocking solution at 4°C. Sections were rinsed three times for 10 min. with PBST and incubated with secondary antibodies (Table 1) diluted in PBST, supplemented with 1 μg/mL of DAPI (Invitrogen, Cat# 1306) at RT for 2 h. Sections were washed three times with PBST, and slides mounted with Mowiol 4-88 (Sigma-Aldrich, Cat# 81381) prepared in 0.1M tris-glycerol. Imaging was done using a Zeiss LSM-800 confocal at 10/20/40X magnifications.

**Table 1.** Primary and secondary antibodies

| Antigen | Primary Ab | Source (Cat. #) | RRID | Dilution | Secondary Ab | Source | Dilution |
| --- | --- | --- | --- | --- | --- | --- | --- |
| AC3 | Rabbit Poly. | ThermoFisher (PA5-35382) | AB_2552692 | 1:300 | Goat anti-Rb IgG Alexa 555 | Invitrogen | 1:1000 |
| α-synuclein endo (M, H) | Mouse IgG1 | BD Bioscience (610786) | AB_398107 | 1:500 (IHC) | Goat anti-Ms IgG1 Alexa 555 | Invitrogen | 1:1000 |
|  |  |  |  | 1:1000 (WB) | Goat anti-Ms IgG1 IRDye 800 | LICOR | 1:10000 |
| α-synuclein (H) | Mouse IgG1 | Millipore Sigma (MABN824) | AB_2921328 | 1:1000 | Goat anti-Ms IgG1 Alexa 488 | Invitrogen | 1:1000 |
| α-synuclein, aggregated | Mouse IgG1 | BioLegend (847801) | AB_2632811 | 1:500 | Goat anti-Ms IgG1 Alexa 488 | Invitrogen | 1:1000 |
| α-synuclein, aggregated | Mouse IgG1k | EMD Millipore (MABN389) | AB_2716647 | 1:300 | Goat anti-Ms IgG1 Alexa 488 | Invitrogen | 1:1000 |
| α-synuclein, aggregated | Rat IgG2a, k | BioLegend (864901) | AB_2810763 | 1:300 | Goat anti-Rat IgG Alexa 546<br>Donkey anti-Rat IgG Alexa 647 | Invitrogen | 1:1000 |
| α-synuclein pSer129 | Mouse IgG2a | BioLegend (825702) | AB_2734593 | 1:1000 | Goat anti-Ms IgG2a Alexa 488 | Invitrogen | 1:1000 |
| α-synuclein pSer129 | Rabbit IgG | Abcam (ab51253) | AB_86997 | 1:500 | Goat anti-Rb IgG Alexa 488 | Invitrogen | 1:1000 |
| β-actin | Mouse IgG1 | GeneTex (GTX629630) | AB_2728646 | 1:1000 | Goat anti-Ms IgG1 IRDye 800 | LICOR | 1:10000 |
| BrdU | Rat IgG | Abcam (ab6326) | AB_305426 | 1:300 | Goat anti-Rat IgG Alexa 488 | Invitrogen | 1:1000 |
| Caspase 3 cleavage | Rabbit Poly. | Cell Signaling (9661) | AB_2341188 | 1:500 | Goat anti-Rb IgG Alexa 546 | Invitrogen | 1:1000 |
| c-Src/Src | Mouse IgG2a | Santa Cruz (sc-5266) | AB_627308 | 1:250 | Goat anti-Ms IgG2a Alexa 555 | Invitrogen | 1:1000 |
| Dlx2 | Rabbit Poly. | EMD Millipore (AB5726) | AB_2093141 | 1:200 | Goat anti-Rb IgG Alexa 555 | Invitrogen | 1:1000 |
| Doublecortin (DCX) | Guinea Pig | EMD Millipore (AB2253) | AB_1586992 | 1:500 | Goat anti-Rb IgG Alexa 555 | Invitrogen | 1:1000 |
| GAP43 | Mouse IgG2a | ThermoFisher (33-5000) | AB_2533123 | 1:100 | Goat anti-Rb IgG Alexa 488 | Invitrogen | 1:1000 |
| GFAP | Rabbit Poly. | BioLegend (840001) | AB_2565444 | 1:1000 | Goat anti-Rb IgG Alexa 555 | Invitrogen | 1:1000 |
| Iba1 | Rabbit Poly. | Wako (016-20001) | AB_839506 | 1:200 | Goat anti-Rb IgG Alexa 555 | Invitrogen | 1:1000 |
| NQO1 | Goat IgG | Abcam (ab2346) | AB_302995 | 1:100 | Donkey anti-Goat Alexa 555 | Invitrogen | 1:1000 |
| OMP | Goat Poly. | Wako (019-22291) | AB_664696 | 1:500 | Donkey anti-Goat Alexa 546 | Invitrogen | 1:1000 |
| Phospho-S6 (pS6) | Rabbit IgG | Cell Signaling (4858) | AB_916156 | 1:400 | Goat anti-Rb IgG Alexa 555 | Invitrogen | 1:1000 |
| TBR1 | Rabbit Poly. | Abcam (ab31940) | AB_2200219 | 1:500 | Goat anti-Rb IgG Alexa 546/647 | Invitrogen | 1:1000 |
| TBR2 | Rabbit Poly. | Abcam (ab23345) | AB_778267 | 1:250 | Goat anti-Rb IgG Alexa 546 | Invitrogen | 1:1000 |
| TH (clone LNC1) | Mouse IgG1 | EMD Millipore (MAB318) | AB_2201528 | 1:500 | Goat anti-Rb IgG Alexa 546 | Invitrogen | 1:1000 |
| VGLUT2 | Rabbit Poly. | SySy (135-403) | AB_887883 | 1:200 | Donkey anti-Rb IgG Alexa 488 | Invitrogen | 1:1000 |

### Electron microscopy

For electron microscopy (EM) analysis, we used 12-14 months WT and α-syn-Tg mice (n=3). Briefly, mice were administered an overdose of Euthasol (Virbac) and transcardially perfused with saline solution followed by 1 ml/g body weight of iced cooled 4% paraformaldehyde (PFA) + 0.2% glutaraldehyde in phosphate buffered saline (PBS) to fix the tissues. After overnight post-fixation at 4°C, 50 μm-thick coronal sections were obtained in a vibratome (Pelco 101). Next, sections were treated with 4% osmium (Electron Microscopy Sciences, 19160), dehydrated with graded alcohols, and stained with uranyl acetate (Electron Microscopy Sciences, 22400) and lead citrate (Electron Microscopy Sciences). For micro-dissection, sections were embedded in EMbed 812 resin (Electron Microscopy Sciences, 14900), and both the OB and anterior piriform cortex (APC) were sectioned at 0.04 mm in an ultramicrotome (UltraCut-E). Microsections were mounted on formvar-coated slot grids (Electron Microscopy Sciences) and images were taken using a JEOL electron microscope (JEM-1200 EX II) at the level of the glomeruli in the OB, and layer I in APC with 12,000X magnification.

### BrdU injections and detection

Neurogenesis in the OB was studied using the thymidine analog 5-bromo-2’-deoxyuridine (BrdU) (52), prepared freshly before injections by diluting powdered BrdU (RPI, Cat# B74200) in sterile 1X-Dulbecco’s PBS solution (Gibco, Cat# 14190-144) at 37°C for 5 min. to obtain a 5 mg/mL solution. Then, mice from the early (6-7 months, n=5/genotype) and late (12-14 months, n=3/genotype) experimental groups received two intraperitoneal injections of 50 mg/kg of BrdU, separated by 2 hours between 1-3 pm to account for circadian variation.

At 25 days post-injection (DPI), mice were perfused, brains cryoprotected and tissues sectioned for IHC (see above). To detect BrdU, sections were thawed at 60°C for 10 min., rehydrated in PBS, and the antigen unmasking was made simultaneously to the DNA denaturalization by incubating sections in a 0.02M HCl at 65°C for 35 min. to break the DNA, followed by an incubation in the same solution in ice for 5 min. Next, acid was neutralized by incubating the sections in 0.1M boric acid-borax buffer (pH 8.6) for 10 min. at RT. The subsequent steps including blocking solution and primary/secondary antibodies incubations were performed identically as described above.

### Proteomics by LFQ-MS

Tissues were obtained from OB and APC of WT and α-syn-Tg mice at 6-7 (n= 3/genotype) and 12-14 (n=3/genotype) months, respectively. Briefly, mice were euthanized with an overdose of 5% isoflurane (Covetrus) and brains were rapidly extracted from skulls. Working on an ice-cooled petri dish, the left and right OBs and APCs were dissected from each animal using a scalpel and a 45° curved forceps (Dumont #5/45). Tissues were immediately frozen in liquid nitrogen and tissues from the left hemisphere were used for proteomic analysis. LFQ-MS was performed at the Yale Mass Spectrometry & Proteomics Resource of the W.M. Keck Foundation Biotechnology Resource Laboratory. Samples were analyzed as technical triplicates. The raw mass spectrometry data will be publicly available upon publication in the PRIDE depository. The data was normalized to internal controls and total spectral counts. Proteins with two or more unique peptide counts were listed using UniProt nomenclature and included for further analysis. A 1.25-fold change and a p-value difference of <0.05 between WT and α-syn-Tg mice were considered as significant. Heat maps for significantly changed proteins were produced using Qlucore Omics Explorer. Volcano plots were produced using GraphPad Prism 9.2 software. Gene ontology (GO) enrichment analysis (53–55) was used to determine significantly affected biological pathways. Pathways with a false discovery rate of < 0.05 were considered as significantly enriched. Graphs depicting significantly affected pathways were produced using GraphPad Prism.

### Western Blots

Western blots were performed using tissues from the right hemisphere obtained as described above. SDS-PAGE and western blots were performed using standard protocols, and images were collected using a LI-COR Odyssey imaging system. Densitometry values were collected using ImageJ. Western blot signal quantifications are represented as mean ± SEM of the endogenous/human α-synuclein, and aggregated α-synuclein (Table 1) detected signal. As loading control, we labeled membranes against beta-actin (Table 1).

### Quantifications and statistical analyses

Data from olfactory tests were first used to independently compare latency times between males and females in the WT and α-syn-Tg groups, and the data were analyzed using an unpaired non-parametric Mann-Whitney U test. Latency data between WT and α-syn-Tg groups were compared separately for the early (6-7 months) and late (12-14 months) groups using an unpaired non-parametric Mann-Whitney U test. For easier representation of the data in graphical form, latency time data were normalized between 0 (minimum latency time = 0 sec.) and 1 (maximum latency time = 300 sec.).

α-Syn pathology along the olfactory pathway was quantified by measuring the stained area for aggregated α-syn and pSer129-α-syn. Both markers were automatically quantified in the OB layers, the *inner cell zone* in the AON and in PC layer 2 by using Fiji (ImageJ) software with the MorphoLibJ integrated library and plugin (56). We first isolated the region of interest (ROI) on a 16-bit maximum projection of a 159.72 x 159.72 μm confocal image of DAPI nuclear staining with a freehand drawing tool. We used DAPI channel to prevent bias when drawing the outline of the layers, and a mask of the drawn ROI was then applied to the corresponding channels stained with antibodies to quantify stained objects within this ROI. The uneven illumination was corrected by applying a “rolling ball” algorithm with a radius of 100 pixels. To address the noise of the aggregated α-syn and pSer129-α-syn channels, we applied the “remove outliers” function with a pixel radius of 2 and threshold of 50. To segment aggregated α-syn and pSer129-α-syn stained pixels from background, we used the Fiji automatic MaxEntropy threshold function. Due to the fainter staining within OB sections, we used the Fiji automatic “Triangle” threshold function in these sections instead. After making a mask of the images, the percent area occupied by aggregated α-syn and pSer129-α-syn segmented objects within the drawn ROI was quantified by applying the “Analyze Particles” function with a minimum object area of 5 pixels. The stained area for both aggregated α-syn and pSer129-α-syn were compared using unpaired t-tests.

From the EM study, we quantified the numbers of mitochondria and synapses (total, asymmetric, symmetric, and reciprocal) from images taken in the two regions where most of the synapses between projection neurons occur, the glomeruli in the OB and layer 1 of APC. Comparisons between WT and α-syn-Tg mice were made using unpaired t-tests.

Dopaminergic cells were quantified on sections stained for the enzyme tyrosine hydroxylase (TH) from 6-7 months mice. The number of TH-positive cells were counted in both the lateral and medial regions of three consecutive sections of the OB (n= 3) in the GL, and numbers averaged. Comparisons between WT and α-syn-Tg were made using an unpaired t-test.

Phosphorylated S6 ribosomal protein (pS6) intensity of staining was quantified in OB mitral cells of 12-14 months mice (n=4/genotype). Images were taken using an Olympus BX51 microscope, and intensity of the pixels was measured in two-line profiles per image, in two images per animal of both lateral and medial regions of the OB. Pixel intensity was measured using the Olympus cellSens Standard 3.1 software. Background line profiles were used to subtract the value from the stained profiles. Values obtained from line profiles per section and regions of the same animal were averaged to get the pS6 pixel intensity for each animal. Statistical comparisons were made using an unpaired t-test.

BrdU^+^ cells were quantified using an Olympus BX51 microscope. The analyzed sections were taken from midway through the OB and were selected by finding the sections from each animal at the same coronal plane. The right OB sections were chosen arbitrarily, with left OB sections used at planes where the right section was torn or otherwise unusable. BrdU^+^ cells were manually counted via fluorescence microscopy at 20X magnification. The area of the glomerular layer (GL), granular cell layer (GCL) and bulbar rostral migratory stream (OB-RMS), were calculated by manual tracing of an image of the section taken at 4X magnification. Cell counts were divided by their respective areas to calculate average count per area, then averaged over the sections to find the average count per area in a select region, over a given animal. Statistics for BrdU numbers on the average counts per area in the GL, GCL and the OB-RMS were compared separately between early (6-7 months) and late (12-14 months) stages using unpaired t-tests.

Western blot quantifications for human/mouse endogenous α-syn and aggregated α-syn were normalized against the respective beta-actin quantifications. Comparisons between groups using this normalized data were made using unpaired t-tests.

All statistical analysis were carried out using the GraphPad Prism 9.2 software.

## Results

### Olfactory deficits found in late-stage α-syn-Tg mice

A progressive loss of olfaction is a well-known non-motor symptom in PD patients (4, 57). Here, we evaluated olfactory deficits in a mouse model of PD before and after the appearance of motor abnormalities, i.e., early (6-7 months) and late (12-14 months) stages, respectively (45, 58). Olfactory deficits were analyzed using standardized buried food tests as they are effective methods to evaluate the ability of trained mice to locate scented foods using olfactory cues (48, 49, 51).

Our results from olfactory tests showed that α-syn-Tg mice displayed olfactory deficits only at late stages (12-14 months) compared to WT controls (p=0.029) but not at early stages (6-7 months) (p= 0.8048) (Fig. 1A). These data suggested an olfactory impairment caused by α-syn-Tg overexpression in mice at advanced stages of the disease. In both α-syn-Tg and WT groups, data from males and females were merged after we ruled out the influence of sex on the results. To exclude the influence of motor deficits or lack of motivation by food in mice, particularly at late stages, we performed a surface food test in which the latency to find the food was recorded as a measure of mobility. There were no differences between WT and α-syn-Tg at early (p=0.9709) and late (p=0.1156) stages (Fig. 1B), indicating that motor agility did not influence olfactory performance, even though 12-14 months mice had motor deficits on more challenging motor tests (data not shown; (45)). In summary, we found that aged α-syn-Tg mice exhibited olfactory behavior deficits.

**Figure 1.**
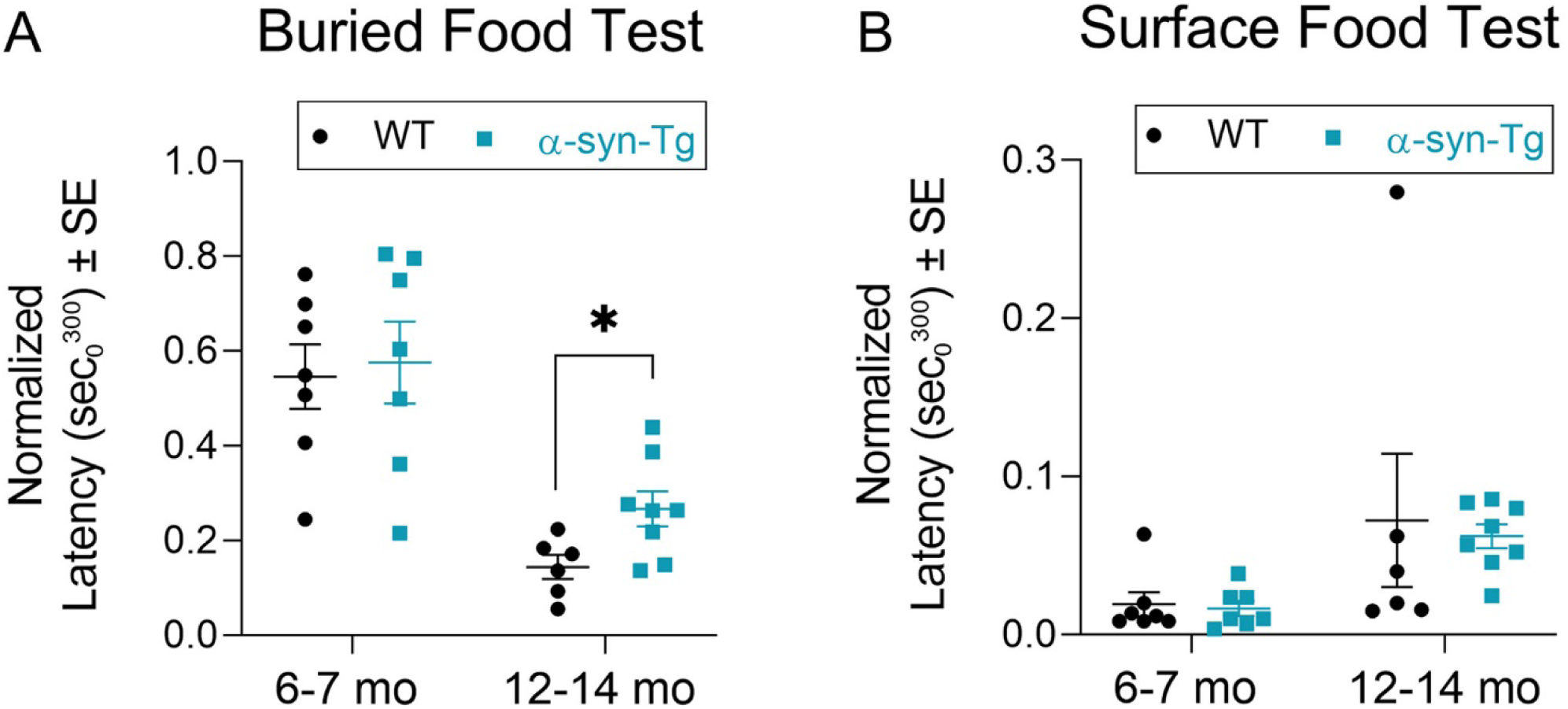
α-syn-Tg mice show olfactory deficits at 12-14 months. (A) Wild type (WT) and α-syn-Tg (carrying the hA30P mutant α-syn) mice are evaluated for olfactory deficits using standard buried food tests. Mice at early stages of the disease (6-7 months) do not exhibit olfactory dysfunction measured as the latency to find a hidden food. At late stages (12-14 months), α-syn-Tg mice show olfactory deficits indicative of dysregulated olfactory pathway. (B) Surface food location test did not show any difference between WT and α-syn-Tg mice, performed to exclude bias due to motor deficits or lack of motivation for food. Data are normalized between 0 as the minimum latency time (0 sec), and 1 as the maximum latency time (300 sec) (* = p<0.05). Statistics: Unpaired non-parametric Mann-Whitney U test.

### Cytoarchitecture of the olfactory pathway of α-syn-Tg mice

Next, we sought to establish any α-syn pathology within the primary olfactory system to investigate the causes of olfactory deficits observed in 12-14 months mice. The olfactory pathway is organized in three fundamental processing stations interconnected by projection neurons (Fig. 2A):

1. Olfactory sensory neurons (OSNs) expressing odor receptors are located in the olfactory epithelium (OE) lining the nasal cavity. When odorants bind, the information is sent to the first olfactory processing center of the brain, the olfactory bulb (OB).
2. The OB has a laminar organization formed by 6 layers which go from the most superficial to the deepest as follows: (1) olfactory nerve layer; (2) glomerular layer (GL); (3) external plexiform layer (EPL); (4) mitral cell layer (MCL); (5) internal plexiform layer (IPL); and (6) granule cell layer (GCL). These layers are formed by a variety of projection neurons (PNs), involved in receiving and processing the olfactory signals received from OSNs, and interneurons (INs) that modulate those signals. PNs include mitral and tufted cells (M/Tc), whose cell bodies locate in the MCL and outer EPL, respectively. PNs extend both primary apical dendrites that arborize in spherical structures called glomeruli (Gl) where they form synapses with axon termini of OSNs, and lateral secondary dendrites that extend laterally within the EPL, where olfactory information is highly modulated. Both type of PNs send their axons to higher brain structures, predominantly the tubular striatum (TuS) and the olfactory cortex (OC), the last the subject of our analysis.
3. The OC is the third station studied in this work and is divided in the anterior olfactory nucleus (AON) and piriform cortex (PC), both responsible for the higher brain processing of an odor and memory formation (59).

**Figure 2.**
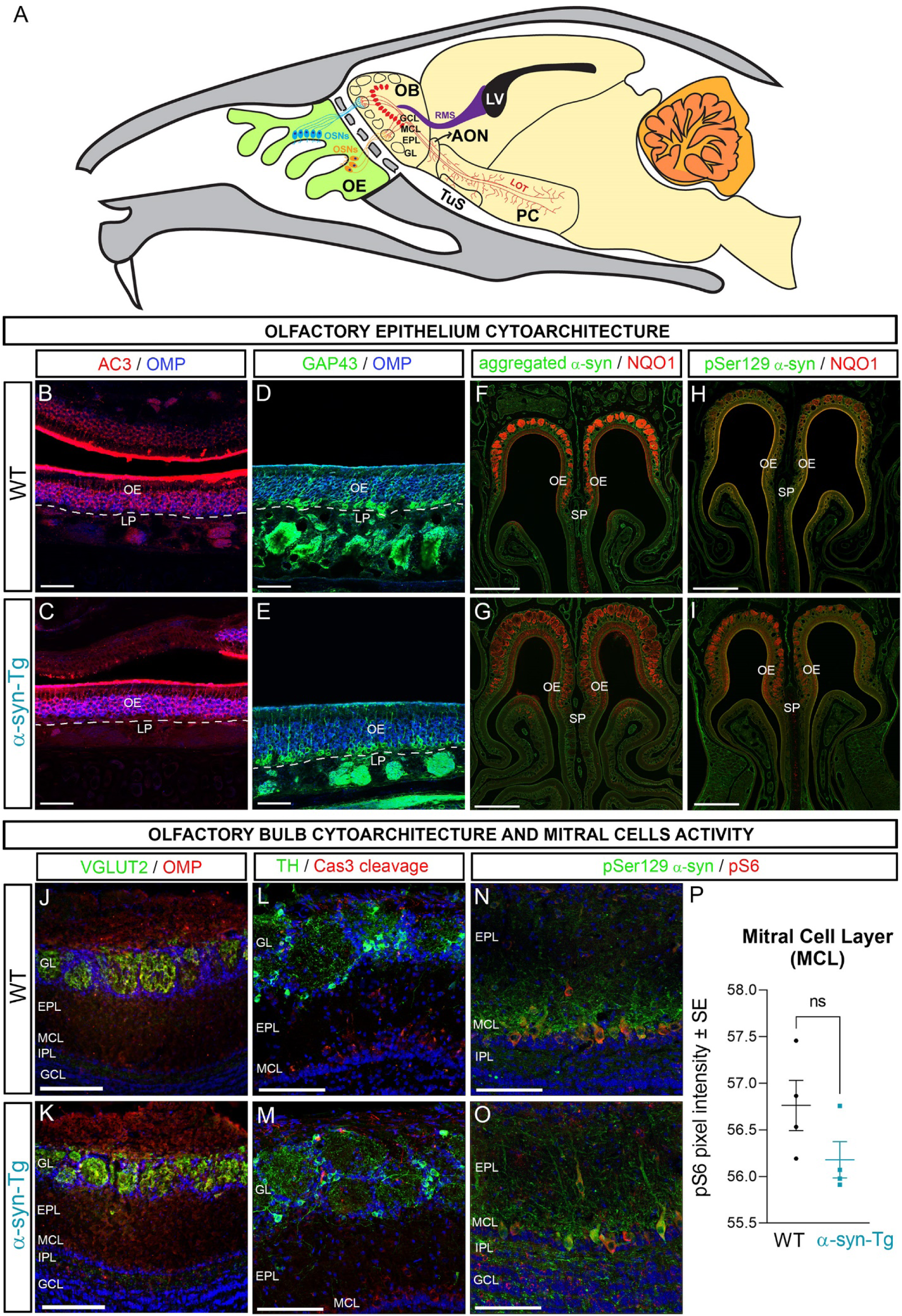
Organization of mouse olfactory system, and cytoarchitecture of the olfactory epithelium (OE) and bulb (OB) in α-syn-Tg mice at 12-14 months. (A) Diagram of the mouse olfactory system summarizing the different station relays used by the olfactory information to travel from the OE to the brain. (B, C) Immunohistochemistry against AC3 (red) and OMP (blue) to label intermediate differentiating and mature OSNs, respectively. (D, E) Staining for GAP43 (green) and OMP (blue) showing immature and mature OSNs, respectively. No alterations in the OE cytoarchitecture are seen in α-syn-Tg mice. (F, G) Low magnification images of the OE stained for α-syn pathology using aggregated α-syn (green; Mouse EMD Millipore, Table 1) and NQO1 (red), specific for dorsal OE. No evidence of α-syn aggregation is seen in the OE. (H, I) Low magnification images of the OE stained for α-syn pathology using pSer129-α-syn (green, Rabbit Abcam, Table 1) and NQO1 (red). No evidence of α-syn pathological phosphorylation is seen in the OE. Differences in NQO1 staining correspond to slight variations in the section plane along the rostral-caudal axis between the different mice. All these staining show an unaltered cytoarchitecture in the OE of α-syn-Tg mice with lack of α-syn pathology. (J, K) OSNs axons labeled with VGLUT2 (green) and OMP (red) in the OB show no differences in the glomerular structure between WT and α-syn-Tg mice. (L, M) Staining of dopaminergic neurons with TH in the OB-GL (green) and signs of apoptosis using Caspase 3 cleavage (red), showing no disruptions in the morphology or cellular processes or increased cell death in α-syn-Tg mice. (N, O) Staining for α-syn pathology in the OB-MCL using pSer129 α-syn (green; Mouse BioLegend, Table 1) and cellular activity detected using pS6 (red) in mitral cells. (P) Quantification of pS6 expression levels shows no significant differences between WT and α-syn-Tg mice. Nuclei counterstained with Dapi (blue). Abbreviations: AC3: adenylyl cyclase type 3; AON: anterior olfactory nucleus; EPL: external plexiform layer; GAP43: growth associated protein-43; GL: glomerular layer; GCL: granule cell layer; LOT: lateral olfactory tract; LP: lamina propia; LV: lateral ventricle; MCL: mitral cell layer; NQO1: NAD(P)H dehydrogenase [quinone] 1; OB: olfactory bulb; OE: olfactory epithelium; OMP: olfactory marker protein; OSNs: olfactory sensory neurons; PC: piriform cortex; pS6: phosphorylated S6 Ribosomal Protein (Ser235/236); RMS: rostral migratory stream; SP: septum; TuS: tubular striatum; TH: tyrosine hydroxylase; VGLUT2: vesicular glutamate transporter 2. Scale bars of 500 μm in F-I; 200 μm in J, K; 100 μm in L, M; 50 μm in B-E, N, O. Statistics: Unpaired t-test.

We began by examining disruptions in the OE neuronal cytoarchitecture. OSNs in the OE are continuously renewed during the lifetime of an individual in a process that requires cell division of OE neural stem cells (horizontal and globose cells) and their differentiation to OSNs. The differentiation stages of OSNs were identified using specific markers such as (1) olfactory marker protein (OMP) to label mature OSNs, (2) adenylate cyclase 3 (AC3) to label differentiating OSNs, and (3) growth associated protein 43 (GAP43) to label immature OSNs (60). Our analysis showed no disrupted OE cytoarchitecture between WT controls (Fig. 2B, D) and α-syn-Tg mice at 12-14 months of age (Fig. 2C, E). We tested if the unaffected morphology was due to the absence of the disease in the OE by detecting two hallmarks of α-syn pathology, specifically: (1) the accumulation of α-syn aggregates (aggregated α-syn); and (2) phosphorylation of serine 129 of α-syn (pSer129-α-syn) (20, 61). The data showed that α-syn pathology was not detected in the OE, and there was no difference between genotypes in aggregated α-syn (Fig. 2F, G) and pSer129-α-syn (Fig. 2H, I) staining. We also assessed for changes along the dorsal-ventral axis by staining with NADPH dehydrogenase [quinone] 1 (NQO1), a selective marker for the dorsal region, and we did not find any difference between the groups (Fig. 2F-I). Our observations were consistent with data found from PD patients (62).

Next, we explored the anatomy of the OB layers and general activity of OB projection neurons in 12-14 months mice. We first studied the GL, formed by the glomeruli and periglomerular neurons (PGNs) by staining against vesicular glutamate transporter 2 (VGLUT2) and OMP, both characteristic markers for OSNs axons. No evidence of anatomical disruptions was found in α-syn-Tg mice compared to WT controls (Fig. 2J, K). In addition, we studied the dopaminergic local circuits established by PGNs expressing the enzyme tyrosine hydroxylase (TH^+^) (63), by quantifying TH^+^ cells around randomly selected glomeruli. Our results showed similar morphologies (Fig. 2L, M) and no statistical differences (p=0.1358; t=1.864, df=4) in the number of TH^+^ cells in WT (443±43 cells/mm^2^) versus α-syn-Tg mice (535±74 cells/mm^2^). These data differ from those previously published showing a decrease in the number of TH^+^ cells, or an increase in the TH innervated area in transgenic mice expressing hA30P mutant α-syn (44, 64, 65). Finally, we searched for changes in the cellular activity in mitral cells as the main projection neurons of the OB, by analyzing the expression levels of phosphorylated S6 ribosomal protein (Ser235/236) (pS6). Oscillations in pS6 levels have been correlated with the activation of neurons in response to stimuli, synaptic plasticity or pathological events such as traumatic injury or seizures (66). In the OB, pS6 expression levels are reported to be reduced in OB mitral cells after sensory deprivation (67). Here, we quantified the intensity of pS6 staining along the MCL in both WT and α-syn-Tg mice, showing no statistically significant difference between the groups (Fig. 2N-P). Overall, these results suggested no effect of α-syn pathology on the gross cytoarchitecture and mitral cell activity of the OB in aged α-syn-Tg mice.

### Accumulation of α-synuclein pathology in α-syn-Tg mice

The α-syn-Tg mouse model of PD expresses the human A30P mutant α-syn under the Thy1 promoter, which is known to trigger protein expression predominantly in projection neurons (68). To validate how this model was affecting the olfactory system, we first investigated where human α-syn (hα-syn) was expressed. We performed triple IHC to detect hα-syn in combination with pSer129-α-syn and aggregated α-syn, as hallmarks of α-syn pathology, in α-syn-Tg mice at both 6-7 and 12-14 months (Fig. 3). We performed the study in two main areas of the olfactory system, the OB and the anterior region of piriform cortex (APC) which is the area that receives the most innervation from the OB (69).

**Figure 3.**
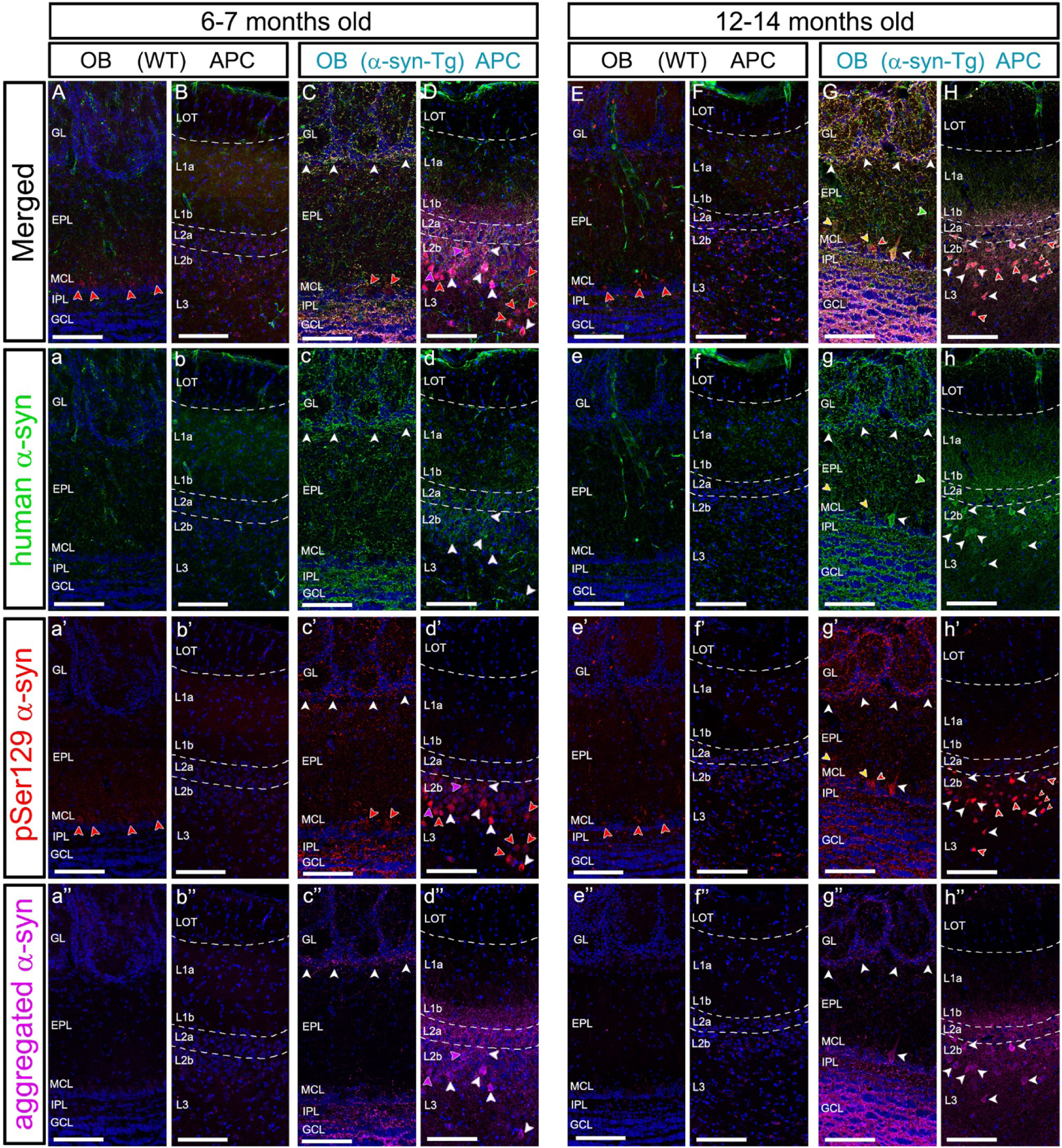
Detection of transgenic human α-synuclein expression and associated α-syn pathology in α-syn-Tg mice. A human specific anti-α-syn antibody (hα-syn) is used to detect the expression of the transgene in the α-syn-Tg mice (green). α-Syn pathology is detected using pSer129-α-syn (red; Rabbit Abcam, Table 1; largely somatic) and aggregated α-syn (magenta; Rat BioLegend, Table 1; largely synaptic). (A-a’’; B-b’’) Staining in the OB and APC of WT control mice at 6-7 months. Only a residual expression of pSer129-α-syn is seen in OB mitral cells (A,a’, red arrowheads). (C-c’’; D-d’’) Staining in the OB and APC of α-syn-Tg mice at 6-7 months. Transgene expression in the OB is seen in neuronal processes of the GL, EPL, IPL and GCL layers. (c). In APC, hα-syn stains neuronal processes of layers 1, 2 and 3 and it appears somatic in some neurons of 2b (d). Most of these processes and neuronal bodies in APC co-expressed α-syn pathology (white arrowheads) (hα-syn^+^; red arrowheads in C, c’, D, d’; magenta arrowheads in D, d’, d’’). Strong expression of aggregated α-syn is detected in layers 1b and 2a of APC (d’’). (E-e’’; F’f’’) Staining in the OB and APC of WT control mice at 12-14 months. A residual expression of pSer129-α-syn remains in the OB mitral cells at this age (E, e’, red arrowheads). (G-g’’; H-h’’) Staining in the OB and APC of α-syn-Tg mice at 12-14 months. Compared with 6-7 moths, at 12-14 months there is a stronger accumulation of the transgene (hα-syn^+^, green) along all layers of OB (g), and layers 1b, 2a and 2b in APC (h). hα-syn staining is somatic in some mitral cells (g). In APC, most neurons expressing hα-syn also show pathology (H-h’’). In APC no hα-syn or α-syn pathology is seen in layer 2a (H-h’’). Nuclei counterstained with Dapi (blue). Abbreviations: APC: anterior piriform cortex; EPL: external plexiform layer; GCL: granule cell layer; GL: glomerular layer; IPL: internal plexiform layer; LOT: lateral olfactory tract; MCL: mitral cell layer; OB: olfactory bulb. Scales of 100 μm.

As expected, hα-syn was not detected in WT animals (Fig. 3A-a, B-b, E-e, F-f), although we observed some residual expression of pSer129-α-syn in the OB-MCL (Fig. 3a’, e’) that did not occur in APC (Fig. 3b’, f’). In transgenic mice, the hα-syn expression pattern was synaptic and appeared distributed predominantly in neuronal fibers in the GCL and IPL of the OB at 6-7 months (Fig. 3c, white arrowheads). However, the hα-syn staining pattern became more somatic in the MCL at 12-14 months (Fig. 3g, white arrowheads). In APC, the hα-syn expression pattern was found predominantly in, 1) the densely packed cell layers 2a+b and 3 formed by superficial and deep pyramidal neurons, respectively (Fig. 3d, h), with higher expression at 12-14 months (compare Fig. 3d staining with 3h). In layers 2 and 3 a more pronounced somatic expression was observed at 12-14 months (Fig. 3d, h, white arrowheads); 2) in layer 1 stronger staining was evident at 12-14 months, suggesting an increase of hα-syn expression with age (Fig. 3D-d, H-h). Sublayer 1a contains the primary synapses from the OB projection neurons onto the apical dendrites of the pyramidal neurons in layers 2 and 3 while sublayer 1b includes the associational synapses with cortico-cortical axons coming from other OC regions (69). α-Syn pathology was detected in both OB and APC tissues from mice at 6-7 (Fig. 3C-c’’, D-d’’) and 12-14 (Fig. 3G-g’’, H-h’’) months, colocalizing predominantly with cell bodies and processes expressing hα-syn (white arrowheads). The overall increase of total α-syn (both human and mouse endogenous) and aggregated α-syn in α-syn-Tg mice was confirmed by WB in OB homogenates of α-syn-Tg mice when compared to WT at both 6-7- and 12-14-months (Fig. 4A, B). The data were not significant at 6-7 months probably because of the insensitivity of the antibody in western blotting (Fig. 4B).

**Figure 4.**
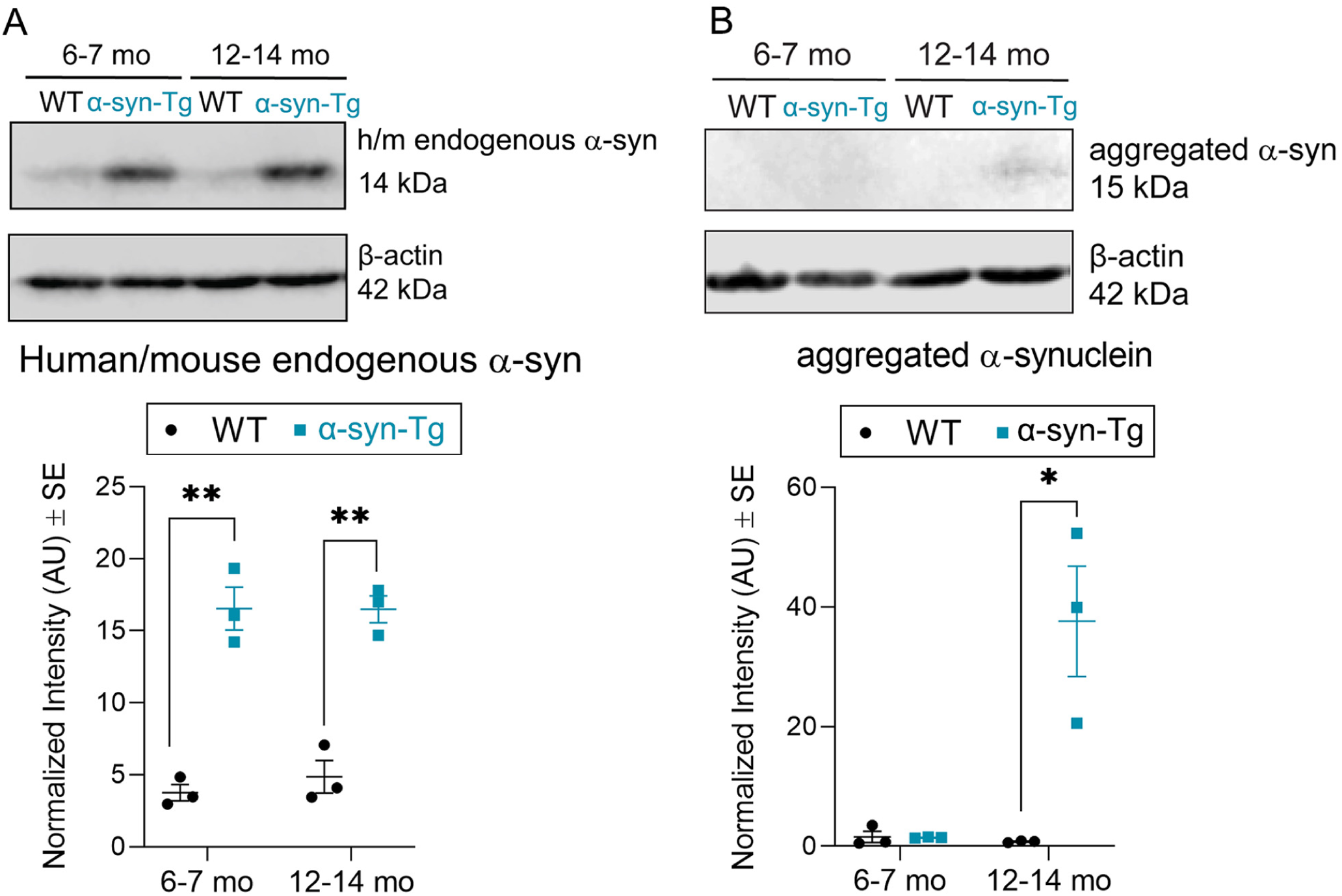
Quantification of human/endogenous and aggregated α-synuclein by western blot (WB). (A) WB detecting total (endogenous and human) α-syn in WT controls and α-syn-Tg mice at both studied stages (6-7- and 12-14-months mice). The amount of protein quantified by densitometry showed a significant increase of total α-syn between WT vs α-syn-Tg mice at both 6-7 and 12-14 months in the OB (B) Detection of aggregated α-syn (Mouse EMD Millipore, Table 1) by WB showed an increase in the amount of protein in the OB of α-syn-Tg mice at 12-14 months β-actin is used as a loading control. *= p<0.05, **= p<0.01. Statistics: Unpaired t-test.

To further investigate which neuronal types were targeted by the overexpression of hA30P mutant α-syn, we performed a triple IHC analysis along the olfactory pathway of 12-14 months mice, to detect hα-syn together with aggregated α-syn as a marker for pathology and either TBR1 (T-Box Brain Protein-1) to identify excitatory/glutamatergic projection neurons (70), or GAD65/67 (Glutamic Acid Decarboxylase isoform 65 and 67) to identify GABA inhibitory neurons (71) (Fig. 5). Our results with TBR1 showed that all cells expressing hα-syn colocalized with TBR1 in the OB-MCL, AONl and APC (Fig. 5B-b’, C’, c’, D’, d’, white arrowheads) but not in the OB-GL region (Fig. 5A-a’). However, not all TBR1 cells expressed hα-syn (Fig. 5A-D, red arrowheads, and red cells). α-Syn pathology studied with aggregated α-syn colocalized predominately with TBR1 cells that also expressed hα-syn (Fig. 5a’’-d’’, white arrowheads). There were instances of GAD65/67 overlapping with hα-syn or aggregated α-syn (Fig. 5E-h’’, red arrowheads), but it was not as robust as seen in projection neurons and was less somatic.

**Figure 5.**
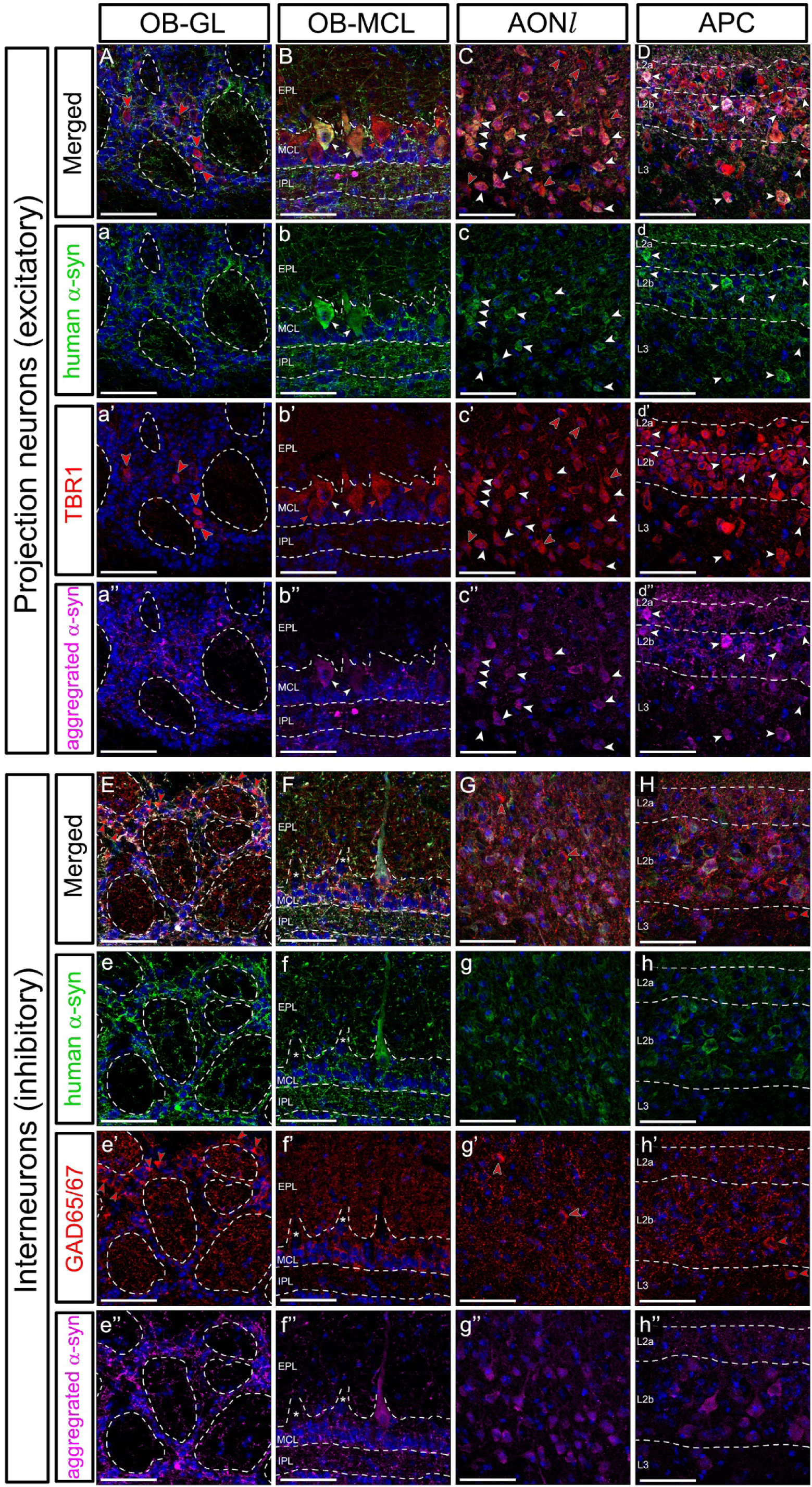
Detection of the transgenic human α-synuclein expression in excitatory projection and inhibitory neurons at 12-14 months. (A-a’’ to D-d’’) Staining for human specific α-syn (hα-syn; green) with the projection neurons marker TBR1 (red) and aggregated α-syn (magenta, Rat BioLegend, Table 1) in the OB: GL and MCL layers, the lateral AON and APC. No colocalization is seen in TBR1^+^ neurons in the OB-GL with hα-syn or aggregated α-syn (A-a’, red arrowheads). In the OB-MCL, AONl and APC, some neurons express the transgene (hα-syn^+^) which always colocalized with TBR1 and α-syn pathology (B-b’’, C-c’’, D-d’’, white arrowheads). (E-e’’ to H-h’’) Staining for human specific α-syn (hα-syn; green) with the inhibitory neurons marker GAD65/67 (red) and aggregated α-syn (magenta, rat BioLegend, Table 1) in the same olfactory regions. GAD65/67^+^ neuron colocalizes with hα-syn or aggregated α-syn. The IPL, which is occupied predominantly by OB projection neuron axons is labeled for hα-syn or aggregated α-syn but not for GAD65/67, indicating that the transgene is prominent in projection excitatory neurons. Nuclei counterstained with Dapi (blue). Abbreviations: AONl: lateral anterior olfactory nucleus; APC: anterior piriform cortex; EPL: external plexiform layer; GAD65/67: glutamate decarboxylase isoforms 65 and 67; IPL: internal plexiform layer; MCL: mitral cell layer; OB: olfactory bulb; TBR1: T-Box Brain Transcription Factor-1. Scales of 50 μm.

Collectively, we concluded that the α-syn pathology in the α-syn-Tg mouse model was found throughout the olfactory pathway, most prominently in projection/excitatory neurons.

### α-Synuclein pathology in the central olfactory pathway of α-syn-Tg mice

We next investigated α-syn pathology in the three main stations of the central olfactory pathway: the OB, the AON, and PC of mice at 12-14 months. In the OB, we quantified the expression of both aggregated α-syn and pSer129-α-syn in each of the OB layers and found that α-syn pathology was laminar specific (Fig. 6). There was no difference in GL expression between WT and α-syn-Tg mice in either aggregated α-syn (Fig. 6C; p=0.7187, t=0.3866, df=4) or pSer129-α-syn (Fig. 6J; p=0.4241; t=0.8893, df=6). In the EPL, minimal expression of aggregated α-syn was seen in both WT and α-syn-Tg groups (Fig. 6A-a’’, B-b’’, D; p= 0.7187; t=0.3866, df=4). In WT and α-syn-Tg mice pSer129-α-syn staining was equivalent in the EPL (Fig 6H-h’’, I-i’’, K; p=0.2336; t=1.402, df=4). A subpopulation of OB projection neuron somata expressing TBR1^+^ in the outer EPL (oEPL), tufted cells, were pSer129-α-syn labeled in α-syn-Tg mice (Fig. 6h, h’).

**Figure 6.**
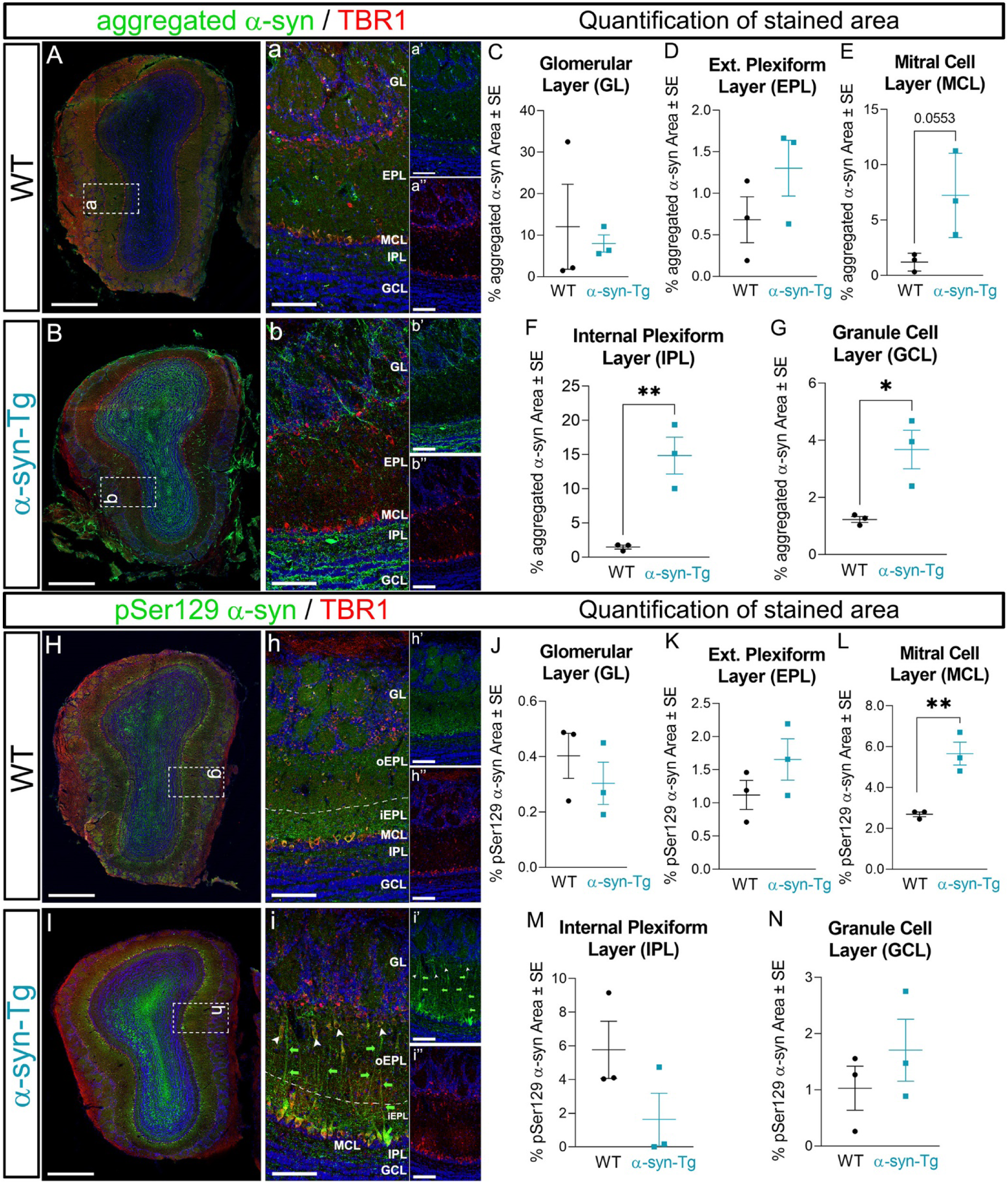
α-Synuclein pathology in the olfactory bulb (OB) of mice at 12-14 months. (A-a’’, B-b’’) Representative images of OB coronal sections of mice stained against aggregated α-syn (green, Mouse BioLegend, Table 1) and TBR1 (red) as a marker for projection neurons in WT (A-a’’) and α-syn-Tg (B-b’’) mice. Staining seems to be restricted to cell processes. Quantification of stained area shows a significantly higher accumulation of aggregated α-syn in α-syn-Tg mice in the IPL (F) and GCL (G), with a strong trend in MCL (E). No differences are detected between the groups in GL (C) and EPL (D). (H-h’’, I-i’’) Representative images of OB coronal sections of mice stained against pSer129-α-syn (green, Mouse BioLegend, Table 1) and TBR1 (red) in WT (H-h’’) and α-syn-Tg (I-i’’) mice. Quantifications show significantly increased pSer129-α-syn in the MCL of α-syn-Tg mice compared to controls (L) with no statistical differences in the other layers (J, K, M, N). Remarkably, a strong accumulation of pSer129-α-syn is seen in tufted cells (i, i’, white arrowheads) and in apical and lateral dendrites of mitral cells (i, i’, green arrowheads) of α-syn-Tg. There is a characteristic staining of pSer129-α-syn in the GCL forming a pattern intermingled with granule cell islets that likely belong to M/Tc axons (I). Nuclei counterstained with Dapi (blue). Abbreviations: EPL: external plexiform layer; GCL: granule cell layer; IPL: internal plexiform layer; M/Tc: mitral and tufted cells; TBR1: T-Box Brain Transcription Factor-1. Scale bars of 500 μm in A, B, H, I; 100 μm in a-a’’, b-b’’, h-h’’, i-i’’. *= p<0.05; **= p< 0.01. Statistics: Unpaired t-test.

The lateral dendrites of mitral cells extend horizontally in the inner EPL (iEPL) and synapse with granule cell dendrites, while apical dendrites extend radially in the EPL to innervate glomeruli. Both dendrites were stained with pSer129-α-syn in WT (Fig. 6h, h’) and α-syn-Tg mice (Fig. 6i, i’, green arrows), but the distribution was different between the experimental groups: the expression was limited to the lateral dendrites in the WT mice while both the lateral and apical dendrites showed expression in the α-syn-Tg mice (Fig. 6h, i). As mentioned above, we attributed its presence in the WT mice with a residual phosphorylation of α-syn previously reported to occur in healthy aged, adult brains representing ~4% of the total brain phosphorylated α-syn (20). In the MCL, there was a strong trend of higher levels of aggregated α-syn in α-syn-Tg mice (Fig. 6E; p=0.0553; t=2.679, df=4), while pSer129-α-syn expression was significantly higher in α-syn-Tg mice compared to WT (Fig. 6L; p<0.01; t=5.204, df= 4). Labeling of pSer129-α-syn which was most evident in the cell bodies of mitral cells of both WT (Fig. 6H-h’’) and α-syn-Tg (Fig. 6I-i’’) mice, colocalized with TBR1. These data together with our previous observations using hα-syn (Fig. 5A-d’’), were consistent with the notion that α-syn pathology, detected by phosphorylation of α-syn in Ser129, is a process restricted to projection/non-GABAergic neurons, as previously reported (72). Deeper in the IPL, aggregated α-syn was significantly higher in α-syn-Tg (Fig. 6b, b’) than in WT controls (Fig. 6F; p<0.01; t=4.934, df=4), while pSer129-α-syn expression levels were unchanged between experimental groups (Fig. 6M; p=0.1465; t=1.798, df=4). This is consistent with the observation that pSer129-α-syn staining is mainly concentrated in cell bodies and thicker neurites (73). In the GCL aggregated α-syn (Fig. 6A-a’’, B-b’’) expression was significantly higher in α-syn-Tg than in WT (Fig. 6G; p<0.01; t=3.594, df=4) and appeared preferentially localized to neurites interdigitated with the granule cell islets. The staining for pSer129-α-syn within the GCL (Fig. 6H-h’’, I-i’’) showed a pattern characteristic of the mitral/tufted cell (M/Tc) axons as they coalesce and subsequently form the lateral olfactory tract (LOT). There was only a trend toward a higher expression of pSer129-α-syn in the GCL (Fig. 6N; p=0.3740; t=0.998, df=4). Together, our results indicated that α-synuclein pathology accumulation was enhanced in the OB projection M/Tc neurons of α-syn-Tg mice.

We next examined the olfactory cortices formed by the anterior olfactory nucleus (AON) and piriform cortex (PC) for evidence of α-syn pathology. The AON is formed by a compact ring of cells, the *pars principalis,* surrounding the olfactory peduncle. This region is subdivided into two basic zones: an *outer plexiform layer (opl)*, and an *inner cell zone (icz)* containing pyramidal cells (74). We found a trend for increased aggregated α-syn in the *inner cell zone* of α-syn-Tg mice (Fig. 7A-C; p=0.1033; t=2.103, df=4). Interestingly, pSer129-α-syn was significantly increased in projection neurons (TBR1^+^ cells colocalizing with pSer129-α-syn) of the AON *inner cell zone* in α-syn-Tg mice with respect to controls (Fig. 7J-K; p<0.01; t=5.852, df=4), consistent with the pathology restricted to these cells. These results showed compelling evidence of AON pathology as described in humans (75, 76).

**Figure 7.**
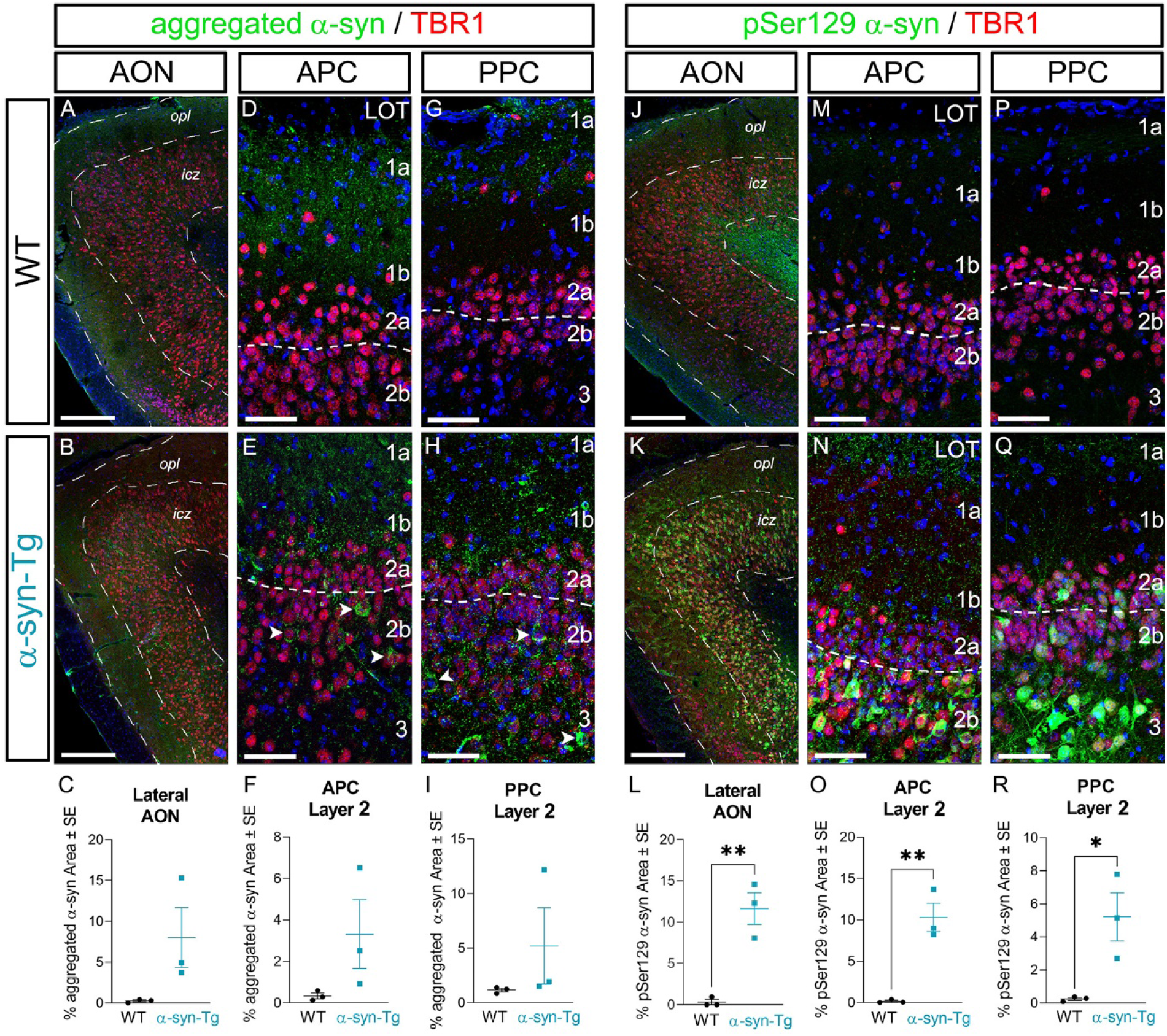
α-synuclein pathology in anterior olfactory nucleus and piriform cortex of mice at 12-14 months. (A, B) Representative images of AON coronal sections from WT and α-syn-Tg mice stained against aggregated α-syn (green, Mouse, BioLegend, Table 1) and TBR1 (red) as a marker for projection neurons. (C) Percentage of stained area in the AON showing higher levels of aggregated α-syn in α-syn-Tg vs WT mice with no statistical differences due to high variability. (D, E, G, H) Representative images of aggregated α-syn staining from the APC and PPC showing α-syn aggregation in layer 2 (a + b) of α-syn-Tg mice, with accumulation in some neuronal cell bodies (arrowheads). (F, I) Quantification of aggregated α-syn stained area in APC and PPC showing no statistical differences between WT and α-syn-Tg mice, although there is a trend to be higher in α-syn-Tg mice. (J, K) Representative images of pSer129-α-syn staining (green, Mouse BioLegend, Table 1) in AON, showing a strong accumulation of α-syn pathology in cell bodies of projection neurons co-labeled with TBR1 (red). (L) The comparison of stained area for pSer129-α-syn in AON shows statistically significant higher accumulation in α-syn-Tg mice. (M, N, P, Q) Examples of images taken in APC and PPC showing an intense labeling for pSer129-α-syn in neuronal cell bodies of layer 2 (a + b) in APC, and IIb in PPC (dotted line). (O, R) Quantification of pSer129-α-syn stained area in APC and PPC showing statistically significant differences between WT and α-syn-Tg mice. Nuclei counterstained with Dapi (blue). Abbreviations: AON: anterior olfactory nucleus; APC: anterior piriform cortex; *icz*: inner cell zone; *opl*: outer plexiform layer; PPC: posterior piriform cortex; TBR1: T-Box Brain Transcription Factor-1. Scale bars of 200 μm in A, B, J, K; 50 μm in D, E, G, H, M, N, P, Q. *= p<0.05; **= p< 0.01. Statistics: Unpaired t-test.

In PC, we studied areas in the anterior- (APC) and posterior- (PPC) piriform cortex to cover the anatomical and functional differences along the rostral-caudal axis (59, 69). In the APC, we found an increase in aggregated α-syn in layers 1b, occupied predominantly by the apical dendrites of PC projection neurons (69) (Fig. 7D, E; compare similar staining on Fig. 3d’’, h’’), and layer 2b, occupied predominantly by pyramidal neuron cell bodies some of which were strongly labeled for aggregated α-syn (Fig. 7E, arrowheads). Quantification of the stained area showed a trend of higher aggregated α-syn in α-syn-Tg mice, although this increase was not statistically significant most likely due to the high variability among individual mice (Fig. 7F; p=0.1489; t=1.785, df=4). Results for aggregated α-syn in PPC were analogous (Fig. 7G-I; p=0.3128; t=1.154, df=4). In contrast, the expression levels of pSer129-α-syn highlighted an intense pathological profile in α-syn-Tg mice in both APC (Fig. 7M-O; p<0.01; t=5.897, df=4) and PPC (Fig. 7P-R; p<0.05; t=3.383, df=4) compared to WT controls. In these cases, pSer129-α-syn staining was restricted to cell bodies and processes of projection neurons in layers 2a and b, and layer 3 (Fig. 7N and Q) of α-syn-Tg mice. Interestingly, there was a lack of pSer129-α-syn labeling in layer 2a of APC compared to PPC (Fig. 5N, Q). Sublayer 2a is occupied by semilunar cells (SL) (77), and the reasons for the differential expression in APC vs. PPC requires further analyses.

Collectively, these data showed substantial evidence of α-syn pathology throughout the central olfactory pathway, with a prominent accumulation in projection neuron cell bodies and processes, whose pathology seems likely to be linked to olfactory deficits at late stages of PD.

### Ultrastructural analysis revealed unaltered synaptic and mitochondrial numbers in α-syn-Tg mice

We next asked if synaptic connectivity was impacted in α-syn-Tg mice at 12-14 months. We used EM to quantify number and type of synapses found in the two primary regions in the olfactory pathway where projection neurons make synaptic contacts: (1) the OB glomerular neuropil (Gl), where M/Tc receive excitatory asymmetrical synapses from OSN axons and inhibitory symmetrical synapses with periglomerular interneurons (78); and (2) layer 1 of the APC, where axons from M/Tc make asymmetrical excitatory synapses with apical dendrites of PC pyramidal cells in layer 1a and local interneurons make symmetrical inhibitory synapses with the PC pyramidal apical dendrites in layer 1b (69, 79). In both regions, we quantified asymmetric (Fig. 8, red arrowheads), symmetric (Fig. 8, green arrowheads), and reciprocal (Fig. 8, yellow arrowheads) synapses. We also quantified the number of mitochondria, whose dysfunction is reported to play a determining role in the progression of PD and can also be used as a proxy for neuronal activity (80).

**Figure 8.**
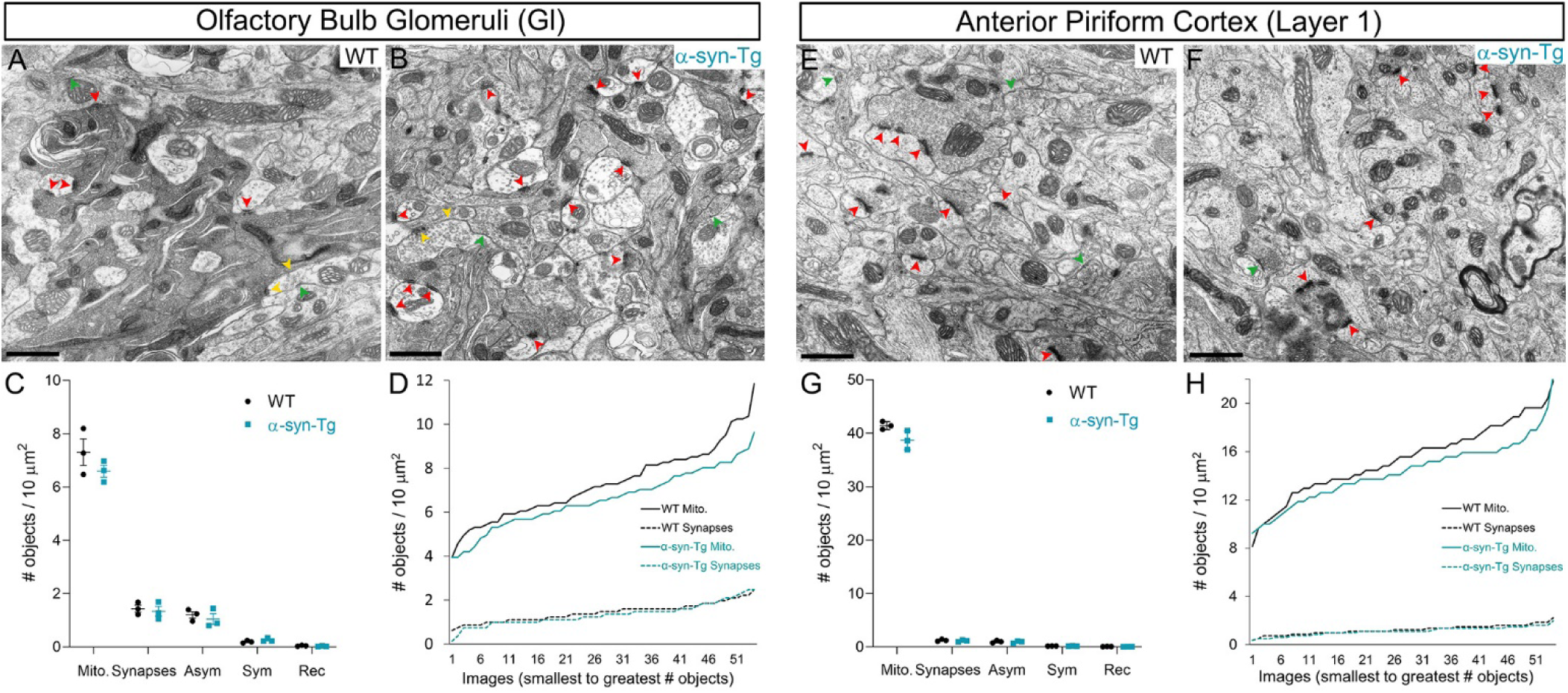
Ultrastructure of synapses and mitochondria in the OB glomeruli and APC layer I. (A, B) Representative images of electron microscopy in the glomeruli of OBs from WT and α-syn-Tg mice. Number of mitochondria (Mito) and different types of synapses are quantified and highlighted as: asymmetric (Asym. Red arrowheads), symmetric (Sym. Green arrowheads) and reciprocal (Rec. Yellow arrowheads). (C) Quantification of Mito. and synapses in the glomeruli parenchyma show no statistical differences between WT and α-syn-Tg mice. (D) Representation of the number of Mito. and total synapses counted per image and arranged from the smallest to the greatest number of objects, showing a strong trend of α-syn-Tg mice to have less Mito. although no statistical differences are found. (E, F) Representative images of APC layer 1 parenchyma at EM showing asymmetric (red arrowheads) and symmetric (green arrowheads) synapses. (G) Quantification of mitochondria and synapses in APC showing no statistical differences between WT and α-syn-Tg mice. (H) Representation of the distribution of mitochondria and synapses on each image arranged from lowest to highest number of objects showing a similar trend to that observed in OB. Scale bars of 0.5 μm. Statistics: Unpaired t-test.

Total numbers of synapses and mitochondria between WT and α-syn-Tg mice in both the Gl (Fig. 8A-C) or APC (Fig. 8E-G) parenchyma showed no statistical differences: number of synapses in OB (p=0.6819; t=0.4412, df=4) and APC (p=0.4953; t=0.7494, df=4), asymmetric synapses in OB (p=0.5449; t=0.6609, df=4) and APC (p=0.7396; t=0.3563, df=4), symmetric synapses in OB (p=0.2292; t=1.418, df=4) and APC (p=0.7096; t=0.400, df=4), reciprocal synapses in OB (p=0.2301; t=1.414, df=4) and APC (p=0.1929; t=1.564, df=4), and mitochondria in OB (p=0.2605; t=1.309, df=4) and APC (p=0.0685; t=2.475, df=4). These data indicated that anomalies in number of mitochondria or synapses were unlikely to account for olfactory dysfunction. Interestingly, when the data is arranged by individual images from lower to highest number of elements (mitochondria or synapses), we found a consistent trend for fewer mitochondria and synapses in α-syn-Tg mice compared to WT controls (Fig. 8D, H). Collectively, our data did not show evidence of morphological and ultrastructural changes that could explain the olfactory deficits observed in α-syn-Tg mice at late stages (12-14 months).

### Adult neurogenesis in the olfactory system of α-syn-Tg mice

Previous studies suggested that overexpression of human WT or hA30P mutant α-syn in animal models of PD results in altered neurogenesis in different layers of the OB (44, 81–83). Neurogenesis of OB interneurons occurs throughout the lifetime of an individual and is crucial to the control and modulation of olfaction (84, 85). Newly generated interneurons proliferate from neural stem cells located in the subventricular zone (SVZ) of the lateral ventricles (LV), after which they migrate towards the OB in a migratory chain called the rostral migratory stream (RMS) (86). New neuroblasts in this pathway take an average of 12-14 days to reach the OB in adult mice (87) and approximately another week to integrate in the OB circuits (88). Here, we sought to evaluate the effect of α-syn pathology on the neurogenesis of interneurons targeted to the OB at early (6-7 months) and late (12-14 months) stages of the disease, that is, prior to and after the onset of hyposmia. Newly formed neurons were identified by injecting BrdU, and tissues analyzed at 25 days post-injection (DPI). We quantified BrdU^+^ cells in three layers of the OB receiving interneurons from this route, namely, the GL, GCL and bulbar part of the RMS.

In the GL, we found no difference in the number of BrdU^+^ periglomerular neurons (PGNs) at 6-7 months by genotype (Fig. 9E; p=0.7213; t=0.3695, df=8) but showed a significant decrease at 12-14 months (Fig. 9E; p=0.0491; t=2.460, df=6), indicating that the hA30P mutant α-syn affected the neurogenesis at late stages of the disease progression (Fig. 9A-D, a’-d’, E). Since our quantification of PGNs that were TH+ showed no differences between WT and α-syn-Tg mice (see above) and considering that no BrdU^+^-PGNs expressed TH (Fig. 10E, F), we concluded that the PGN population affected by the expression of hA30P mutant α-syn at 12-14 months did not include dopaminergic PGNs. In the GCL, we detected a significant decrease in the number of BrdU^+^ cells only at early (6-7 months) stages (Fig. 9F; 6-7 months: p=0.0136; t=3.152, df=8; vs 12-14 months: p=0.095; t=1.976, df=6), highlighting that hA30P mutant α-syn overexpression was affecting OB neurogenesis in the GCL prior to the onset of olfactory deficits (Fig. 9A-D, a-d, F). This is consistent with results from similar animal models expressing the hA30P α-syn mutation under the CaMK-tet regulated promoter (82). Absence of statistical significance at late stages (12-14 months) may be due to the global decrease in neurogenesis of cells targeting the OB during aging (87, 89), which may have masked the effect of hA30P mutant α-syn. Finally, there was no evidence of a change in the number of BrdU^+^ cells in the OB-RMS at either 6-7 (Fig. 9G; p=0.6048; t=0.5386, df=8) or 12-14 months (Fig. 9G; p=0.1076; t=1.890, df=6) (Fig. 9A-D, a-d). Since BrdU injections at adult stages could only label OB interneurons generated in the SVZ and considering that α-syn pathology was only expressed in projection neurons of the olfactory pathway (Figs. 3,5-7), the effects observed in OB neurogenesis are presumably indirect, for instance via an alteration in the dopamine release into the SVZ (90, 91). Collectively, our data showed that hA30P mutant α-syn affected neurogenesis in the GCL of OB at early stage before the onset of olfactory deficits, whereas neurogenesis in the GL of OB is affected in the later stages when olfactory deficit was apparent in α-syn-Tg mice.

**Figure 9.**
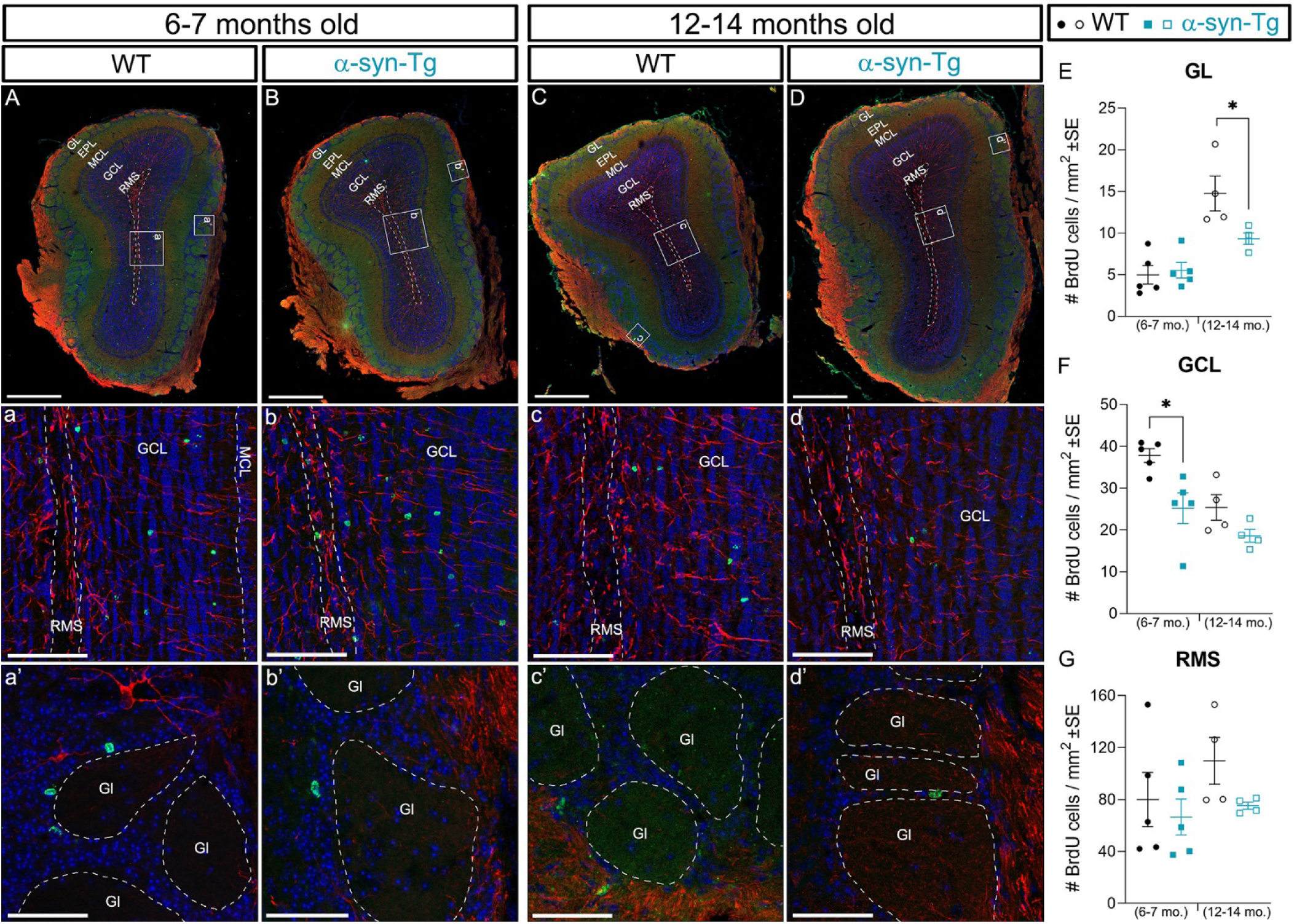
Effect in adult neurogenesis in the olfactory bulb glomerular and granule cell layers of α-syn-Tg mice. Neurogenesis is studied by counting newly generated cells 25 days after they incorporated the thymidine analog BrdU during cell division at the moment of injection. BrdU staining is shown in green, while immature neurons (neuroblasts) are identified with doublecortin (DCX) marker (red). (A to d’) Representative images of OB coronal sections in WT and α-syn-Tg mice at 6-7 months (A-a’, B-b’) and 12-14 months (C-c’, D-d’) stages Insets are high magnification images of BrdU and DCX labeled cells located in the RMS + GCL (a-d) or the GL (a’-d’). (E) Quantification of BrdU cells in the GL shows a statistically significant decrease of cells in α-syn-Tg mice at late stages 12-14 months, and an overall increase in the total number of BrdU^+^ cells in this layer between 6-7 to 12-14 months mice. (F) Analysis of BrdU cells in the GCL shows a statistically significant decrease of labeled cells in α-syn-Tg mice exclusively at early stages of the disease (6-7 months) (G) Quantification of BrdU^+^ cells in the RMS showing no statistical differences between WT and α-syn-Tg mice. Nuclei counterstained with Dapi (blue). Abbreviations: EPL: external plexiform layer; GL: glomerular layer; Gl: individual glomeruli; GCL: granule cell layer; MCL: mitral cell layer; RMS: bulbar portion of the rostral migratory stream. Scale bars of 500 μm in A-D; 100 μm in a-d; 50 μm in a’-d’. *= p<0.05. Statistics: Unpaired t-test.

**Figure 10.**
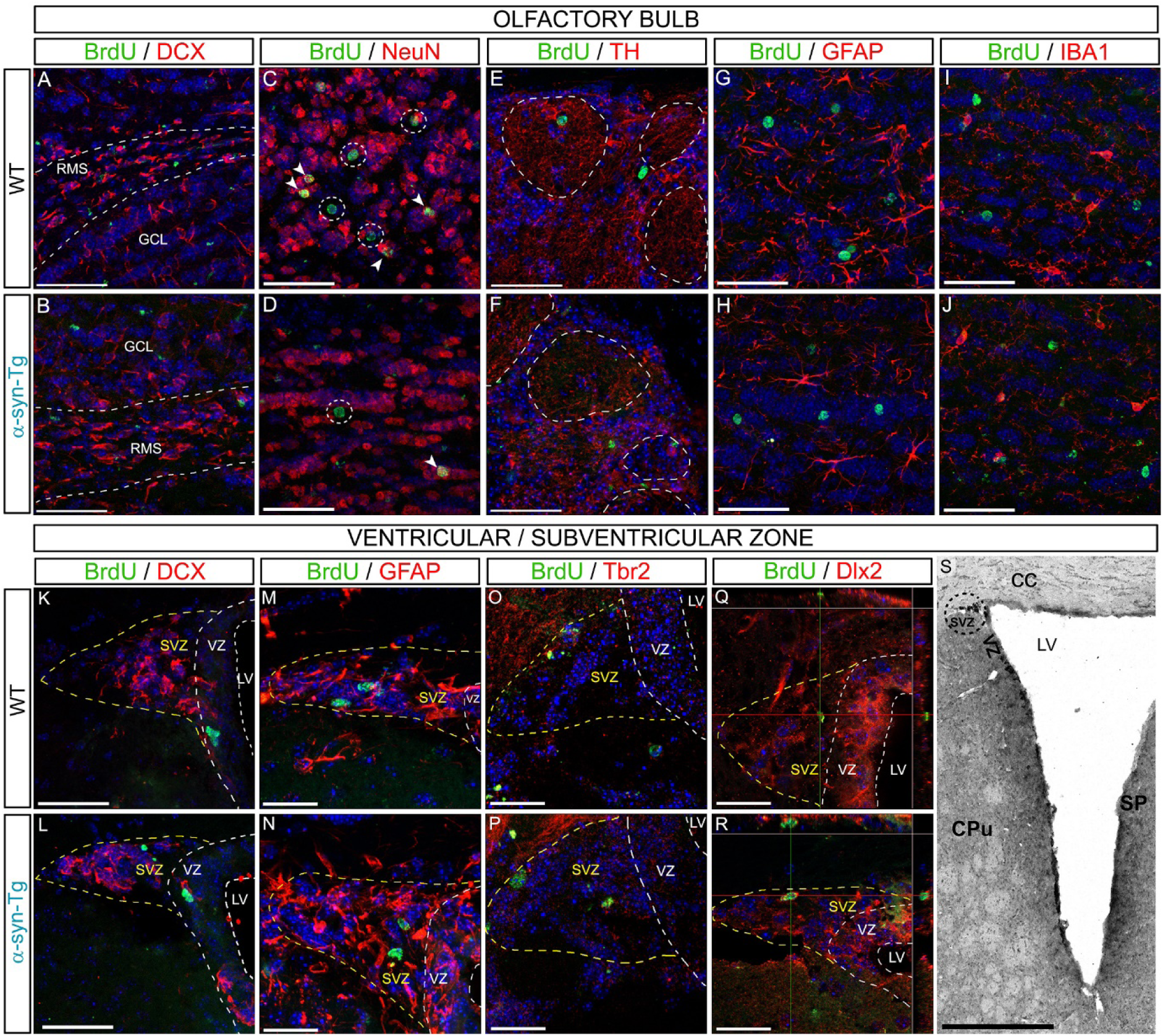
Characterization of cellular phenotypes of BrdU-expressing cells in the olfactory bulb and subventricular zone after 25 days of injection. BrdU staining (green) is shown with different markers typical of neuronal and glial cells (red). (A-J) Phenotypic characterization in the OB. (A, B) no BrdU^+^ cell expresses the neuroblast marker DCX in either the GCL or RMS. (C, D) Identification of mature neurons with NeuN in the GCL, which accounts for roughly 43% and 30% of the BrdU^+^ cells in WT controls and α-syn-Tg mice, respectively. (E, F) Detection of dopaminergic neurons in the GL using the TH markers. No colocalization is seen with BrdU^+^ cells. (G, H) GFAP marker is used to identify astrocytes shows no colocalization with BrdU^+^ cells. (I, J) Microglial cells expressing IBA1 do not colocalize with BrdU^+^ cells either. (K-S) Phenotypic characterization in the SVZ of WT control and α-syn-Tg mice. (K, L) DCX marker is used to label neuroblast or “A” cells which do not colocalizes with BrdU^+^ cells. (M, N) GFAP is used to label radial glia neural stem cells or “B” cells do not colocalizes with BrdU^+^ cells. (O, P) TBR2 used to identify SVZ-dorsal TACs do not colocalizes with BrdU^+^ cells. (Q, R) Dlx2 is used to label TACs or “C” cells which colocalizes with adult resident BrdU^+^ cells at the time of study (25 days post-BrdU injection). (S) Greyscale image of the analyzed region showing the anatomical location of the SVZ next to the ventricular zone (VZ) and the lateral ventricle (LV). Nuclei counterstained with Dapi (blue). Abbreviations: DCX: doublecortin; Dlx2: Distal-Less Homeobox 2; GCL: granule cell layer; GFAP: glial fibrillary acidic protein; IBA1: ionized calcium-binding adapter molecule-1; NeuN: neuronal nuclei marker; OB: olfactory bulb; RMS: rostral migratory stream; SVZ: subventricular zone; TACs: transient amplifying cells; TBR2: T-Box brain transcription factor-2; TH: tyrosine hydroxylase. Scale bars of 400 μm in S; 50 μm in A-J; 25 μm in K-R.

To ensure that all BrdU^+^ cells belonged to neuronal lineages and to exclude cell division by local glial cells, we characterized BrdU^+^ cell phenotypes with IHC using specific markers to detect neuroblasts, mature neurons and glial cells in the two regions we found BrdU labeled cells: the OB and the SVZ. First, we looked for neuroblasts by using doublecortin (DCX), a microtubule-associated protein required for the migration of cells. BrdU^+^ cells that were located in the OB did not colocalize with the DCX marker (Fig. 10A, B), suggesting that most BrdU^+^ cells have completed migration and begun to differentiate into neurons at the time of the study. This is consistent with our prior studies showing that OB neuroblasts require 3 weeks to migrate and differentiate into neurons after generation in the SVZ (87, 88). Mature neurons in the OB were identified using the neuronal nuclei marker NeuN. Unexpectedly, only a fraction of the total number of BrdU^+^ cells (representing approximately 40% of the total) colocalized this marker in the OB (Fig. 10C, D), suggesting that the remaining ~60% of BrdU^+^ cells could be neurons in intermediate stages of differentiation with a DCX^-^/NeuN^-^ phenotype. We also looked for an effect in the TH subpopulation within the GL, a group of mature PGNs reported to be affected in PD patients (92) and in similar animal models (44, 65). Our data demonstrated that none of the BrdU^+^-cells in the GL colocalized with the TH marker (Fig. 10E, F). To ensure that the DCX^-^/NeuN^-^ phenotypes did not belong to dividing glial cells, we searched for astrocytes and microglial cells using glial fibrillary acid protein (GFAP) and ionized calcium-binding adapter molecule 1 (IBA1), respectively. Neither GFAP (Fig. 10G, H) nor IBA1 (Fig. 10I, J) colocalized with BrdU^+^ cells, indicating they were not dividing glial cells. In the SVZ, these markers identify characteristic subpopulations of cells that represent the three differentiation stages of OB interneurons: “A” cells or neuroblasts (DCX^+^); “B” cells or neural stem cells, which share a phenotype with radial glial cells (GFAP^+^); and “C” cells or intermediate transient amplifying cells (TACs) (Dlx2^+^ and Tbr2^+^ for dorsal TACs) (86). Our results showed that BrdU^+^ cells found in the SVZ were negative for DCX (Fig 10K, L), GFAP (Fig. 10M, N) and Tbr2 (T-Box Brain Protein-2) (Fig. 10O, P), indicating they were no “A”, “B” or dorsal “C” cells. However, they expressed Distal-Less Homeobox 2 factor (Dlx2), characteristic of all “C” cells (86), indicating that some “C” (TAC) cells remained in the SVZ 25 days after the BrdU injection. These cellular classifications were uniform for both WT and α-syn-Tg mice.

Collectively, we found that the hA30P mutant α-syn expression significantly reduced OB neurogenesis in the GCL at early stages (6-7 months) and in the GL at later stages (12-14 months), respectively. Among the BrdU ^+^ cells found in the OB, ~40% of them were differentiated to neurons (DCX^-^/NeuN^+^), while the remaining ~60% (DCX^-^/NeuN^-^) remained at intermediate stages of neuronal differentiation. In addition, the scarce number of BrdU^+^ cells found in the SVZ were transient amplifying cells or “C” cells.

### Age-related changes in synaptic endocytic proteins in α-syn-Tg mice

To further understand development of olfactory system pathology in an unbiased manner, we performed proteomic analysis on OB and APC from 6-7 months and 12-14 months WT and α-syn-Tg mice (n=3/genotype with technical triplicates) by label-free quantification mass spectrometry (LFQ-MS). Our proteomic analysis of the OBs from 6-7 months mice identified a total of 3328 proteins, 14 of which were significantly differentially expressed in α-syn-Tg mice (Fig. 11A, B. Table 2). Consistent with transgenic overexpression and immunohistochemical results, we observed a significant increase in α-syn expression in the OB of α-syn-Tg mice (SYUA, Fig. 8A, B). This was confirmed via Western Blotting (Fig. 4A). Protein kinase c-Src was the most significantly upregulated protein in the OB of α-syn-Tg mice (SRC, Fig. 11A, B, fold change of 2.88). Among other functions, protein kinase c-Src phosphorylates components of endocytic trafficking (93) and notably, α-syn (94). GO enrichment analysis performed on the proteins that showed significant changes (p>0.05) identified pathways involving vesicle-mediated transport, specifically endocytosis, which were significantly enriched (Fig. 11C). The vast majority of proteins from our dataset that were implicated in these pathways were downregulated in α-syn-Tg samples (Fig. 11A, B), suggesting a deficit in vesicle-mediated transport in the OB. Overall, proteomic analysis of the OB at early stages revealed dysfunction in endocytosis and vesicular transport.

**Figure 11.**
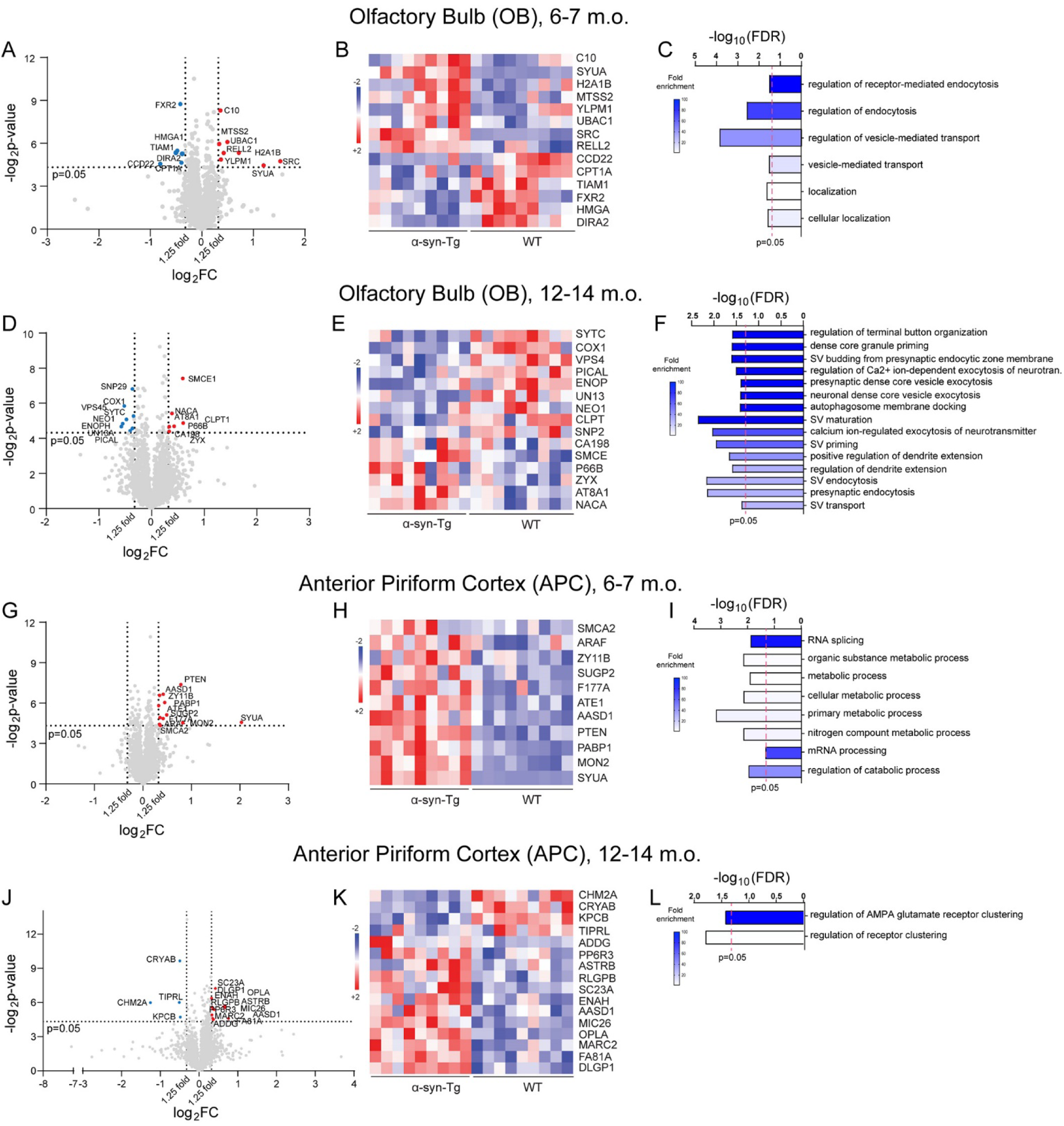
Proteomics of OB and APC revealed defects in synaptic endo- and exocytosis pathways in the OB, but minimum alterations in APC of α-syn-Tg mice. (A) Volcano plot of fold change in the OB proteome of 6-7 months α-syn-Tg compared to control wild type (WT) mice (n=3 mice/genotype). Proteins that changed 1.25-fold (vertical dotted lines) with a p-value of <0.05 (horizontal dotted line) are considered to be significantly differentially expressed. Among the 14 significant proteins, 8 are upregulated (red) and 6 are downregulated (blue). (B) Heat map of significantly differentially expressed proteins in OB of α-syn-Tg compared to WT mice at early stage (6-7 months) for each technical replicate (3 technical replicates/mouse). Red indicates increased expression (+2), and blue indicates decreased expression (−2). (C) Biological pathways that are significantly (FDR<0.05) affected in the OB of 6-7 months α-syn-Tg mice, determined by GO enrichment analysis. (D) Volcano plot of fold change in the OB proteome of 12-14 months α-syn-Tg compared to WT mice (n=3 mice/genotype). Among the 15 significantly differentially expressed proteins (1.25-fold, p<0.05), 7 are upregulated (red) and 8 are downregulated (blue). (E) Heat map of significantly differentially expressed proteins in OB of late stage (12-14 months) α-syn-Tg mice compared to their WT counterparts for each technical replicate (3 technical replicates/mouse). (F) Biological pathways that are significantly (FDR<0.05) affected in the OB of 12-14 months α-syn-Tg mice, determined by GO enrichment analysis. (G) Volcano plot of fold change in the APC proteome. α-syn-Tg compared to WT mice (n=3 mice/genotype) at 6-7 months. All the 11 significantly differentially expressed proteins (1.25-fold, p<0.05) are upregulated in α-syn-Tg mice. (H) Heat map of significantly differentially expressed proteins in APC of α-syn-Tg compared to WT mice at 6-7 months for each technical replicate (3 technical replicates/mouse). (I) Biological pathways that are significantly (FDR<0.05) affected in the APC of 6-7 months α-syn-Tg mice, determined by GO enrichment analysis. (J) Volcano plot of fold change in the APC proteome of α-syn-Tg compared to WT mice (n=3 mice/genotype) at 12-14 months Among the 16 significantly differentially expressed proteins (1.25-fold, p<0.05), 12 are upregulated (red) and 4 are downregulated (blue). (H) Heat map of significantly differentially expressed proteins in APC of α-syn-Tg mice compared to WT controls for each technical replicate (3 technical replicates/mouse) at 12-14 months (I) Biological pathways that are significantly (FDR<0.05) affected in the APC of 12-14 months α-syn-Tg mice, determined by GO enrichment analysis. Abbreviations: APC: anterior piriform cortex; GO: Gene Ontology analysis; OB: olfactory bulb.

**Table 2:** Proteins that are significantly changed in the proteomic analysis of olfactory bulb of α-syn-Tg mice in comparison to WT at 6-7 months of age.

| No. | Accession | Gene name | Protein name | Fold change |
| --- | --- | --- | --- | --- |
| 1 | C10 | Grcc10 | Protein C10 | 1.28436951 |
| 2 | SYUA | Snca | Alpha-synuclein | 2.295515511 |
| 3 | H2A1B | H2ac4 | Histone H2A type 1-B | 1.643448378 |
| 4 | MTSS2 | Mtss2 | Protein MTSS 2 | 1.262605161 |
| 5 | YLPM1 | Ylpm1 | YLP motif-containing protein 1 | 1.292921147 |
| 6 | UBAC1 | Ubac1 | Ubiquitin-associated domain-containing protein 1 | 1.410921697 |
| 7 | SRC | Src | Neuronal proto-oncogene tyrosine-protein kinase<br>Src | 2.876813941 |
| 8 | RELL2 | Rell2 | RELT-like protein 2 | 1.34090359 |
| 9 | CCD22 | Ccdc22 | Coiled-coil domain-containing protein 22 | 0.571394901 |
| 10 | CPT1A | Cpt1a | Carnitine O-palmitoyl transferase 1, liver isoform | 0.759995746 |
| 11 | TIAM1 | Tiam1 | Rho guanine nucleotide exchange factor TIAM1 | 0.719080008 |
| 12 | FXR2 | Fxr2 | Fragile X mental retardation syndrome-related protein 2 | 0.749066485 |
| 13 | HMGA1 | Hmga1 | High mobility group protein HMG-I/HMG-Y | 0.793204904 |
| 14 | DIRA2 | Diras2 | GTP-binding protein Di-Ras2 | 0.770665441 |

In the OB of 12-14 months mice, proteomic analysis detected a total of 2712 proteins, 15 of which were significantly differentially expressed in α-syn-Tg samples (Fig. 11D, E. Table 3). There was a trend in overexpression of α-syn (SYUA), which was confirmed via Western Blot (Fig. 4A). Similar to what we found in 6-7 months mice, GO analysis identified pathways involving vesicle-mediated transport, specifically at the pre-synapses, as significantly enriched (Fig. 8F). These pathways were driven by a majority of proteins that were downregulated in α-syn-Tg samples, including the vesicular trafficking proteins PICALM, UNC13A, SNAP29, and VPS45 (Fig. 11D, E). Thus, proteomic analysis at late stages (12-14 months) reaffirms the endocytic and vesicular transport defects seen at 6-7 months in the OB (Fig. 11A-C), and provided additional evidence that, with age, these deficits became prominent in synaptic terminals of α-syn-Tg mice.

**Table 3:**
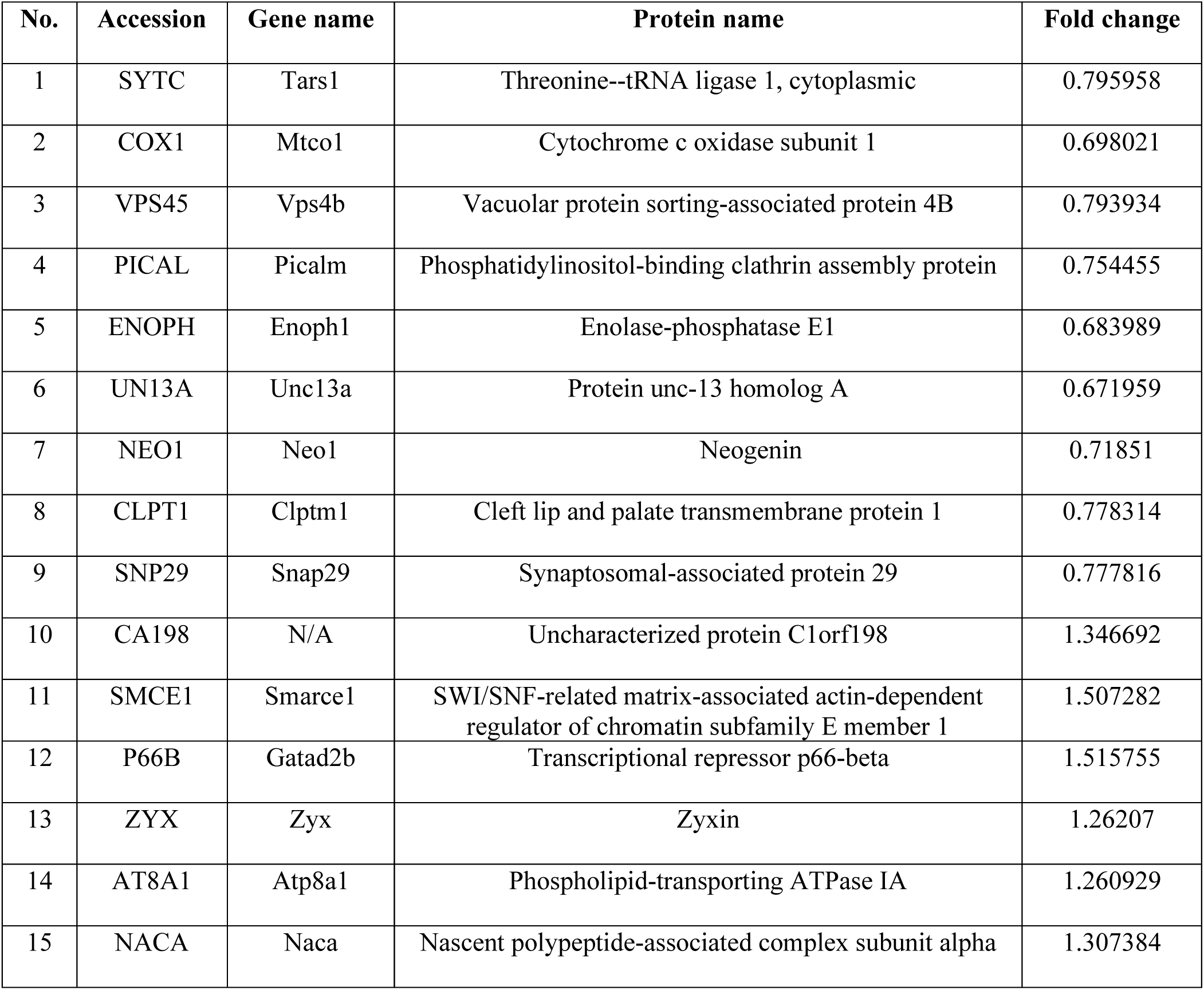
Proteins that are significantly changed in the proteomic analysis of olfactory bulb of α-syn-Tg mice in comparison to WT at 12-14 months of age.

In the APC of 6-7 months mice, proteomic analysis identified a total of 2955 proteins, 11 of which were significantly differentially expressed in α-syn-Tg samples (Fig. 11G, H. Table 4). As anticipated, we observed a significant increase in α-syn expression (SYUA, Fig. 11D, E). Interestingly, GO analysis largely revealed broader dysfunction in cellular metabolism (Fig. 11F) without enrichment of any vesicle mediated-transport pathways seen in the OB (Fig. 11C). In the APC of 12-14 months mice, proteomic analysis identified a total of 2596 proteins, 16 of which were significantly differentially expressed in α-syn-Tg samples (Fig. 11J, K. Table 5). Similarly, GO analysis revealed only a subtle dysregulation in receptor clustering in α-syn-Tg samples, without any dysregulation of synaptic endo- and exocytosis as seen in the OB of these mice. Collectively, our proteomic analysis suggested that the OB is more vulnerable to α-syn pathology induced by the hA30P α-syn mutation expression than the APC, and that olfactory deficits seen at late stages might arise due to early age-related endocytic changes in the synaptic terminals of the OB circuitry.

**Table 4:** Proteins that are significantly changed in the proteomic analysis of anterior piriform cortex of α-syn-Tg mice in comparison to WT at 6-7 months of age.

| No. | Accession | Gene name | Protein name | Fold change |
| --- | --- | --- | --- | --- |
| 1 | SMCA2 | Smarca2 | Probable global transcription activator SNF2L2 | 1.279780958 |
| 2 | ARAF | Araf | Serine/threonine-protein kinase A-Raf | 1.268139535 |
| 3 | ZY11B | Zyg11b | Protein zyg-11 homolog B | 1.268384587 |
| 4 | SUGP2 | Sugp2 | SURP and G-patch domain-containing protein 2 | 1.405654737 |
| 5 | F177A | Fam177a1 | Protein FAM177A1 | 1.27607181 |
| 6 | ATE1 | Ate1 | Arginyl-tRNA--protein transferase 1 | 1.252608747 |
| 7 | AASD1 | Aarsd1 | Alanyl-tRNA editing protein Aarsd1 | 1.335151511 |
| 8 | PTEN | Pten | Phosphatidylinositol 3,4,5-trisphosphate 3-phosphatase and dual-specificity protein phosphatase PTEN | 1.720804806 |
| 9 | PABP1 | Pabpc1 | Polyadenylate-binding protein 1, PABP-1, Poly(A)-binding protein 1 | 1.364366716 |
| 10 | MON2 | Mon2 | Protein MON2 homolog | 1.766115672 |
| 11 | SYUA | Snca | Alpha-synuclein | 4.090515865 |

**Table 5:** Proteins that are significantly changed in the proteomic analysis of anterior piriform cortex of α-syn-Tg mice in comparison to WT at 12-14 months of age.

| No. | Accession | Gene name | Protein name | Fold change |
| --- | --- | --- | --- | --- |
| 1 | CHM2A | Chmp2a | Charged multivesicular body protein 2a | 0.418897 |
| 2 | CRYAB | Cryab | Alpha-crystallin B chain | 0.70911 |
| 3 | KPCB | Prkcb | Protein kinase C beta type | 0.715298 |
| 4 | TIPRL | Tiprl | TIP41-like protein | 0.701951 |
| 5 | ADDG | Add3 | Gamma-adducin | 1.290529 |
| 6 | PP6R3 | Ppp6r3 | Serine/threonine-protein phosphatase 6 regulatory subunit 3 | 1.552139 |
| 7 | ASTRB | Gramd1b | Protein Aster-B | 1.266189 |
| 8 | RLGPB | Ralgapb | Ral GTPase-activating protein subunit beta | 1.260604 |
| 9 | SC23A | Sec23a | Protein transport protein Sec23A | 1.338548 |
| 10 | ENAH | Enah | Protein enabled homolog | 1.293051 |
| 11 | AASD1 | Aarsd1 | Alanyl-tRNA editing protein Aarsd1 | 1.583143 |
| 12 | MIC26 | Apoo | MICOS complex subunit Mic26 | 1.260649 |
| 13 | OPLA | Oplah | 5-oxoprolinase | 1.597227 |
| 14 | MARC2 | Mtarc2 | Mitochondrial amidoxime reducing component 2 | 1.274637 |
| 15 | FA81A | Fam81a | Protein FAM81A | 1.695311 |
| 16 | DLGP1 | Dlgap1 | Disks large-associated protein 1 | 1.309585 |

## Discussion

Olfactory deficits are proposed as prodromal for PD and have the potential to predict the risk and the severity of the disease (6, 95–101). However, the cellular and molecular mechanisms that affect olfaction in PD remain unknown. To study PD, animal models have been developed carrying mutated genes linked with familial forms of PD (39, 102). Here, we used a mouse model of PD carrying the human A30P mutant α-syn expressed under the Thy1 promoter or α-syn-Tg (45) to study the cellular and molecular anomalies associated with olfactory deficits. We focused our analyses on two stages of the disease progression as per the onset of olfactory dysfunction: (1) early-stage mice at 6-7 months, and (2) late-stage mice at 12-14 months.

### Spatiotemporal variabilities of α-synuclein pathology in the olfactory system of α-syn-Tg mice

We screened for hα-syn expression and pathology throughout the olfactory pathway examining the different stations (Fig. 3 and 5). We find broad expression of hα-syn throughout the olfactory pathway with prominent expression in projection neurons (68, 103),. Interestingly, pSer129 α-syn pathology was found mainly in projection neurons, suggesting that these cells drive the olfactory dysfunction seen in these transgenic mice.

The presence of α-syn pathology in the olfactory system of transgenic mice at 6-7 months was unexpected because a behavioral deficit was not detected at the early stage. At advanced stages of the disease progression (12-14 months), the α-syn-Tg mice exhibited significant olfactory deficits in parallel with the α-syn pathology. Why α-syn pathology did not affect the olfactory behavior at early stages remain unknown, but one possibility suggested by our proteomic data was the appearance of changes in the synaptic vesicle trafficking proteome in older mice that were not found in the younger mice. As noted above, the delayed onset of olfactory deficits could be attributed to regulated protein expression controlled by genetic modifiers (104), including those that could act on the gene promoter employed here, Thy1. Some of these modifiers have been reported to interact with the expression of the hA30P mutant α-syn to modify the toxicity of this protein (105).

To establish a mechanism that explained the temporal susceptibility of olfactory neurons to the hA30P mutant α-syn, we investigated the anomalies in synaptic architecture and intracellular signaling pathways in 12-14 months mice. Synaptic transmission is regulated by α-syn through interactions with synaptic vesicle membranes and impacting on the presynaptic architecture (106–108). To explore this in the α-syn-Tg mice, we performed an ultrastructural analysis and quantified excitatory and inhibitory synapses and mitochondria in the projection neurons synaptic regions of the OB and APC, the glomeruli and layer 1, respectively. Surprisingly, these analyses did not show any differences in ultrastructural features that might account for the behavioral deficits in α-syn-Tg mice (Fig. 8). Similarly, although mitochondria function deficits have been reported in PD (Barbuti et al., 2021) we did not find evidence of alterations in the number/density of mitochondria.

We identify two plausible mechanisms to explain the late-stage olfactory deficits in α-syn-Tg mice: (1) decreased neurogenesis; and (2) perturbations of synaptic exo- and endocytosis pathways proteins involved in the synaptic transmission.

### The hA30P mutant α-syn reduces olfactory bulb neurogenesis at both early and late stages of PD

Neurogenesis in the olfactory system occurs throughout the lifetime of an individual, where newly formed neuroblasts join chains of migrating neuroblasts extending to the ependymal core of the OB via the rostral migratory stream (RMS) (85). Once detached from the RMS, neuroblasts migrate radially to their final positions in either the GCL or GL of the OB. In PD, the evidence of aberrant neurogenesis is limited and controversial with some reports showing either decreases in neurogenesis in the SVZ (90), or no effect (109). Here, using BrdU^+^ injections, we found a decrease in OB neurogenesis in α-syn-Tg mice (Fig. 9) that affected primarily the GCL at early (6-7 months) and the GL at later (12-14 months) stages, respectively. A decrease in OB neurogenesis in these two layers was reported previously in similar animal models but at earlier ages (82, 110), so that it is reasonable to speculate on an effect of hA30P mutant α-syn in OB neurogenesis.

Although Zhang et al. (2019) reported cell death in the GCL of a PD mouse model, we did not find any evidence for the staining of cleaved Cas3 in the olfactory pathway (Fig. 2L, M) that would account for the decrease in BrdU^+^ cells. The most plausible explanation for the observed decrease in BrdU^+^ cells is that SVZ proliferation was reduced, perhaps linked to an indirect effect of affected dopaminergic innervation into the SVZ, as reported before (90). Overall, it seems likely that the decrease in neurogenesis, seen as a decrease in BrdU labeling, contributed to the olfactory phenotype we report here. The delayed onset of a clear behavioral olfactory dysfunction suggests strongly that the effects of the α-syn pathology in the olfactory pathway must be cumulative.

### Alterations in OB and APC proteomics of synaptic trafficking in α-synuclein transgenics

We analyzed unbiased proteomics data in WT and α-syn-Tg mice in the OB and APC to determine cellular pathways that were affected by the overexpression of hA30P mutant α-syn. The hA30P α-syn mutation causes α-syn pathology (111, 112) and is implicated in trafficking mechanism including those associated with both endo- and exocytosis (113). The hA30P α-syn mutation was previously associated with disruptions in vesicular trafficking, synaptic dysfunction, apoptosis, and formation of permeable annular pore-like protofibrils in the cell membrane that disrupt transmembrane trafficking (20, 21, 114–116). While 6-7 months mice did not exhibit olfactory deficits, the presence of α-syn pathology in olfactory areas suggested that the underlying mechanisms of olfactory deficits may arise early and could be cumulative leading to the deficits seen at older ages α-syn-Tg mice. Using proteomics at this early stage, we found perturbations of the proteome in both OB and APC brain areas, particularly in those proteins involved in the endocytic and vesicle-mediated transport pathways, which were significantly affected in the OB of α-syn-Tg mice. At later stages when olfactory deficits arose (12-14 months), these defects appear to be exacerbated in the synaptic terminals of the OB. This result was consistent with the fact that mutations in endocytic proteins cause familial PD (117) and that elevated α-syn levels and aggregation can cause inhibition of synaptic vesicle exo- and endocytosis (118–121). Notably, we observed minimal disruption of biological pathways in the APC. These results suggest that the OB is more susceptible than APC to the α-syn pathology and may be a primary site in the olfactory pathway for olfactory behavior deficits seen in α-syn-Tg mice. Thus, together with defective neurogenesis, endocytic dysregulation in the synaptic terminals in the OB might initiate pathological processes leading to olfactory deficits in PD patients, especially those carrying α-syn mutations. Similar mechanisms might also contribute to sporadic PD cases, as majority of them show LB formation and α-syn pathology.

## Conclusions

Our work provides compelling evidence for the following: (1) mice carrying the human A30P mutant α-syn exhibit olfactory deficits at a later stages of the disease progression, when PD-like motor deficits are apparent; (2) transgenic expression causes α-syn pathology along the entire central olfactory pathway, affecting predominantly projection neurons (TBR1^+^); (3) α-syn pathology does not cause structural and morphological abnormalities in the olfactory system; (4) the overexpression of hA30P mutant α-syn causes a decrease in SVZ neurogenesis and interneurons targeted to the OB in the GL and GCL at late and early stages of the disease, respectively; and (5) hA30P mutant α-syn causes decreased expression of proteins involved in synaptic endocytosis and vesicle-mediated transport in the OB of early and late α-syn-Tg mice. This, together with the alterations in neurogenesis, likely explain the olfactory deficits observed at late stages.

## Acknowledgements

We thank Christine Kaliszewski for technical assistance with EM, and John E. Lee for a-syn-Tg and WT animal colonies maintenance and genotyping. We thank Phil Coish for reading the manuscript

